# Fixed-parameter feedback can hold the hand near a target under gravity in two musculoskeletal arm models

**DOI:** 10.64898/2026.06.15.732510

**Authors:** Jun Kobayashi

## Abstract

Can fixed-parameter feedback hold the hand near a target under gravity? Using muscle-length reflex and endpoint feedback in two MyoSuite arm models, we asked whether settings selected with a rigid index finger could still keep its tip near the target when the finger could bend. In the 34-muscle model, we screened 5640 combinations of controller type and parameters across three target arm postures and two update periods. Only 163 combinations (2.9%) met the holding criterion in the final 1.2 s (window-mean hand position within 0.05 m of the target, drift < 0.005 m, maximum joint speed < 0.01 rad/s). Eleven selected settings, including at least one at each target, also passed every evaluation window of 18.6-s trials from all nine initial conditions. When those settings were transferred unchanged to the 63-muscle model, 89 of 99 trials passed the fingertip and shared-joint criteria in all three windows. Only 18 of the same 99 passed the primary criterion, which required all 38 joints to meet the speed limit. In the 71 trials that passed only the shared-joint criterion, middle-finger motion exceeded the speed limit in the first window, but all joints met it in the later windows. At two equilibria in the 34-muscle model, small perturbations grew when feedback was removed and muscle excitation was held constant. Thus a controller can hold the hand near its target over the tested intervals even when the held posture is mechanically unstable under constant excitation.

## 1 Introduction

Holding the hand at a position against gravity is a basic task for a redundant, muscle-driven limb [1]. Whether a given controller accomplishes it in a simulation depends on the controller and on how the periphery is modelled: comparing torque-driven, spring–damper and muscle-based models of fast elbow movements, Pinter et al. found that the robustness of the same movement to perturbations differed between periphery models [2]. In Hill-type muscle models with a serial elastic element, neglecting serial damping can produce high-frequency oscillations [3]. Other simulation studies combined muscle-length (and velocity) feedback with feedforward stimulation in models with compliant tendons: fast single-joint elbow movements [4], perturbed elbow movements [5], spinal posture and flexion under gravity [6], and three-dimensional stance and squatting with a hierarchical PID-based architecture whose lowest level is a muscle-length reflex [7]. Stroeve examined arm posture directly in a neuromusculoskeletal model combining feedforward and feedback control [18]. The equilibrium-point/referent tradition [8] and the internal-model tradition [9, 10] offer different accounts of how posture is held, and whether they compete or complement each other remains debated [11, 12, 13]. A simulation can inform this debate only if it states exactly which plant and which control law were used and how holding was judged.

This paper compares fixed-parameter (time-invariant) controllers for holding the hand near three targets in the 34-muscle MyoSuite myoArm [14, 15]. For reflex and endpoint controllers, we test predefined combinations of parameter values at three target arm postures and two update periods. We assess holding, then test selected settings in longer trials and from perturbed starting states. We test the same combinations in the 63-muscle myoArm, from which the 34-muscle model is derived. Both models use the index fingertip as the endpoint. In the 34-muscle model it is fixed relative to the hand; in the 63-muscle model the index finger can bend and displace the tip even if the arm and wrist keep the same posture. Both criteria assess fingertip holding. The primary criterion also limits speed at all joints, whereas the secondary criterion applies the same joint-speed test in both models. Our main contribution is to identify fixed-parameter feedback controllers that keep the hand near a target under gravity and to document the tested conditions under which they did so in the 34-muscle model. At each of the three targets, at least one target-specific setting met the holding criterion in all three evaluation windows of 18.6-s trials from each of nine initial states. The reflex controllers used muscle-tendon lengths at the target posture as reference values. At each control step, the endpoint controllers used the model to calculate how muscle activation translates into joint torque at the current joint angles and velocities.

Two further findings clarify what such a hold entails.

1. *Hand holding transferred even when the fingers took longer to settle*. With the successful settings transferred unchanged to the 63-muscle model, 89 of 99 trials met the hand and shared-joint criteria. Only 18 met the speed limit at every joint. Middle-finger motion in the first evaluation window accounted for the difference.
2. *Feedback holding can occur at a mechanically unstable equilibrium*. In a post-hoc test at two equilibria reached by successful controllers, some perturbations grew when muscle excitation was held at its equilibrium value. Thus the ability to hold with feedback and the mechanical response to perturbations at fixed excitation can be distinguished in the same settings.

## 2 Methods

### 2.1 Plant

We use the MyoSuite environment `myoArmReachFixed-v0` (MyoSuite 2.12.2, MuJoCo 3.6.0). The parent myoArm combines the MoBL upper-extremity model [16, 17] with MyoHand [14]. The environment builds its model from the 63-muscle myoArm XML by removing the finger joints (the digits become rigid bodies), which leaves 20 hinge joints, 34 muscle actuators and 11 joint equality constraints that implement the shoulder rhythm. The model has nine independent joints (seven arm joints and two thumb carpometacarpal joints) and 11 dependent joints. To define a target configuration, we specify the independent joint angles and compute each dependent joint angle as a polynomial of its corresponding independent joint angle. MuJoCo enforces these polynomial relations with soft equality constraints, so they hold approximately during simulation. The controller sets the muscle excitation *u* ∈ [0, 1]^34^ at each update. Muscle activation *a* is the state that responds to this input and determines muscle force at a given length and velocity. It follows first-order MuJoCo dynamics with activation and deactivation time constants of 10 and 40 ms, respectively. The MuJoCo muscle model computes muscle force by scaling active force with muscle length and velocity and adding passive force. Tendons are inelastic and there is no serial damping element. Joints have the viscous damping specified in the model, 0.05–1.05 N m s/rad. Each joint had an additional rotational inertia of 10^−4^ kg m^2^ (MuJoCo’s `armature` parameter) and zero dry friction. Gravity acts along − *z*. The physics step is *h* = 2 ms (a 1-ms check is reported in Sect. 2.4). We verified that, at fixed joint position and velocity, the generalized actuator force is affine in activation for this model (no force or activation limits), which the allocation below uses. Unless stated otherwise, dimensions in Sects. 2.1–2.4 refer to this 34-muscle model. The 63-muscle myoArm from which it is derived is used in a separate comparison (Sect. 2.5).

### 2.2 Targets, start state and initial conditions

Three hand targets R_low, R_mid and R_high are defined by a joint configuration *q*^*\**^ (Table 1): the dependent joints follow the equality constraints and the target hand position *x*^*\**^ is the forward kinematics of the index-finger-tip site at *q*^*\**^ (Fig. 1a). Each nominal rollout starts at 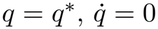. We verified that the simulator started each trial from the prescribed initial state.

**Table 1:** Hand targets. The listed independent joint angles and the default angles of the remaining independent joints define *q*^*\**^ through the equality constraints, which determine the dependent joint angles. The resulting hand position is *x*^*\**^.

| Target | *x*<sup>\*</sup> (*x*, *y*, *z*) (m) | Independent joint angles (rad) |
|---|---|---|
| R_low | (−0.415, −0.344, 0.988) | shoulder_elv 0.5, elbow_flexion 0.7, elv_angle 0.5 |
| R_mid | (−0.415, −0.433, 1.486) | shoulder_elv 1.2, elbow_flexion 1.0, elv_angle 0.6 |
| R_high | (−0.346, −0.296, 1.906) | shoulder_elv 2.0, elbow_flexion 0.6, elv_angle 0.8 |

**Figure 1:**
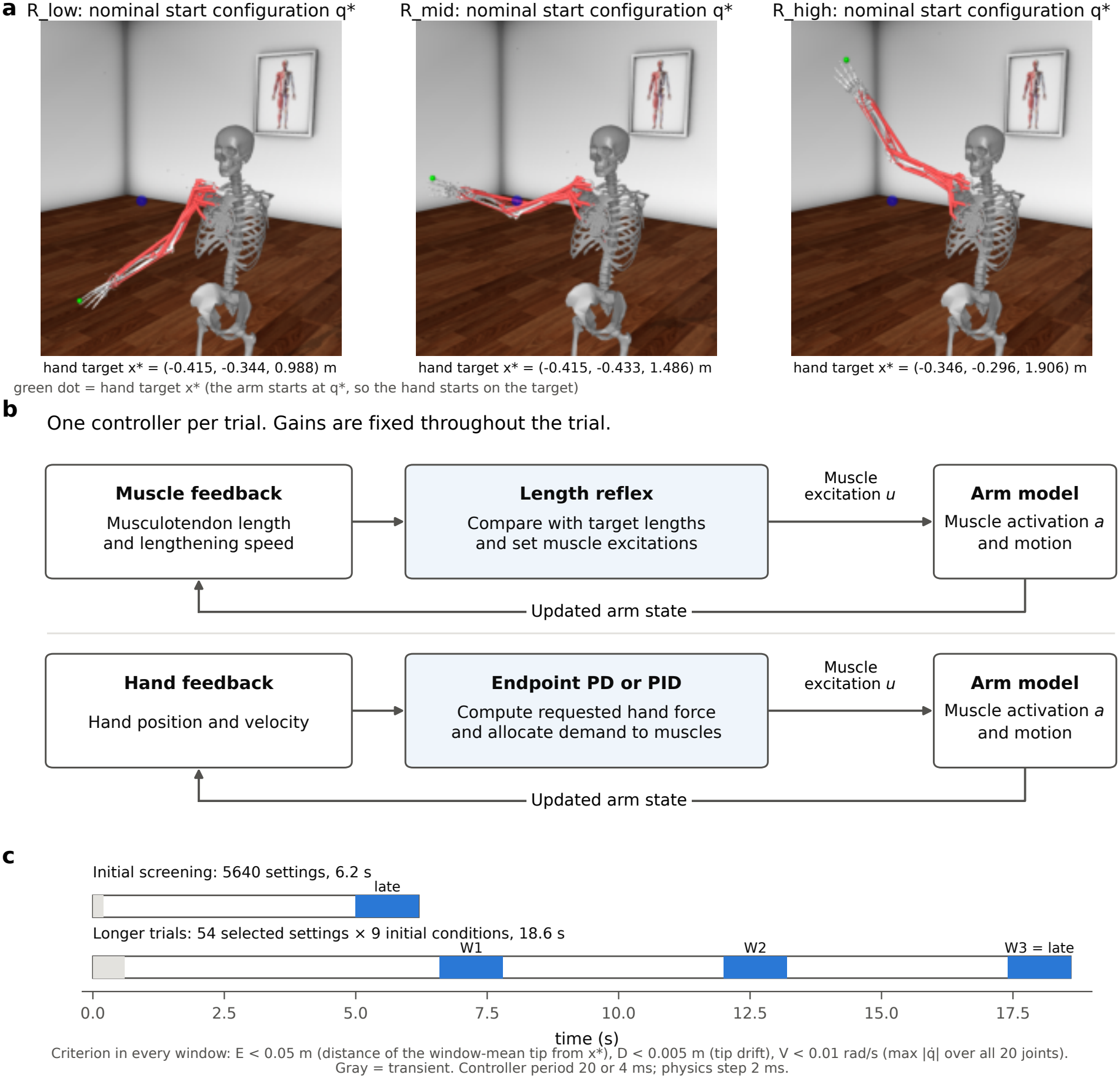
Task, controllers and time design. **a** Nominal start configurations *q*^*\**^ of the 34-muscle model; the green dot is the hand target *x*^*\**^. Perturbed starts add one prescribed single-joint offset. **b** Each trial uses either the reflex or an endpoint PD/PID controller. The reflex uses musculotendon lengths at the nominal target posture as its reference. Endpoint PD/PID converts the requested hand force into a joint torque demand. Depending on the setting, the controller also uses joint feedback for a projected posture correction or includes a gravity-compensation term in the baseline excitation *u*_*b*_. Muscle allocation uses the model’s activation-to-generalized-force map at each update. Both pathways update excitation as the arm moves. **c** The initial screening tests controller settings in 6.2-s trials. The longer trials retest selected settings from nine initial conditions for 18.6 s. Blue bars mark the windows in which the holding criterion is evaluated.

The long-horizon check (Sect. 2.4) adds eight perturbed initial conditions: shoulder elevation or elbow flexion offset by ± 0.02 rad (dependent joints follow the constraints, 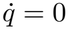), or the velocity of one of these joints set to ± 0.05 rad/s (dependent velocities follow the constraint derivatives). Endpoint controllers can use gravity compensation (GC). For each nominal target posture *q*^*\**^, we use inverse statics to compute the muscle excitation vector *u*_0_ that approximately balances gravity there. The target *x*^*\**^, *q*^*\**^, the reflex reference lengths and *u*_0_ are always those of the nominal target; none is recomputed at the perturbed state.

### 2.3 Feedback controllers for muscle excitation

Each trial uses one of two feedback laws to compute the muscle excitation *u* (Fig. 1b). The muscle-length reflex computes each component *u*_*i*_ directly from feedback about muscle *i*. The endpoint controller uses hand feedback to request a force, then allocates the corresponding torque demand across muscles to obtain *u*. Both controllers are evaluated at a fixed controller period Δ*t* ∈ {20, 4} ms (10 or 2 physics steps), holding *u* constant between control instants. Their parameters do not change during a rollout; the muscle excitation *u* can change, because it depends on the state.

#### 2.3.1 Muscle-length reflex controller

At each update, the reflex controller sets the excitation of each muscle from its own musculotendon length and velocity. For muscle *i*, the reference length *λ*_*i*_ is its musculotendon length at the target posture *q*^*\**^. The controller adds the length deviation *l*_*i*_ − *λ*_*i*_ to a velocity term 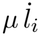, which is positive during lengthening when *µ* > 0. The excitation *u*_*i*_ of muscle *i* is

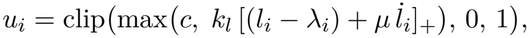

where [*x*]_+_ = max(*x*, 0). The gain *k*_*l*_ (1/m) scales the positive part of the combined signal, and *µ* (s) sets the weight on velocity. The excitation has a minimum value *c* and a maximum of 1. Muscle activation starts at zero.

#### 2.3.2 Endpoint feedback controller (PD or PID)

Following the spring–damper formulation of endpoint impedance [20], the endpoint controller requests a hand force from the position error *e*_*k*_ = *x*^*\**^ − *x*_*k*_ and fingertip velocity 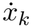at update 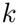:

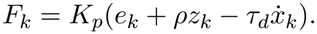

PD omits the integral term. For PID, *z*_*k*_ accumulates fingertip position error over time. Both anti-windup rules cap its contribution; the conditional rule also freezes the integral when accumulation would increase an unrealizable demand. The controller maps the hand force to torques in the independent joint coordinates. Settings with posture feedback add a torque that biases these joints toward the target posture using the kinematic null space of the hand task (*K*_*q*_ = 100 N m/rad, *τ*_*q*_ = 0.1 s).

At each update, bounded least squares allocates the requested torques across muscles using the current activation-to-torque map. Feedback adjusts muscle excitation relative to a fixed baseline *u*_*b*_. With GC, this baseline is the gravity-compensation vector *u*_0_ with a floor of *c* applied to each muscle; initial muscle activation is *u*_0_. Without GC, every muscle has the same baseline *c* and starts with zero activation. The resulting muscle excitations lie between *c* and 1. Allocation assumes instantaneous activation; the simulated muscles follow the 10/40-ms activation dynamics. Section A.2.1 gives the force map, joint projection and allocation equations.

### 2.4 Screening and testing for sustained holding

The experiment had two stages (Fig. 1c). In the initial screening (stage B), we tested each controller setting from its nominal start. In the long-horizon trials, we tested selected settings over a longer interval and from nearby initial states. Both stages used the same holding criterion.

#### Holding criterion

A hold requires the hand to stay near the target, the hand to drift little within the evaluation window, and all joints to move slowly throughout that window. In a window of post-step control samples, let 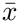 be the mean hand position. The window passes if 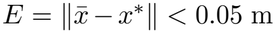, the drift 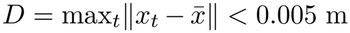, and 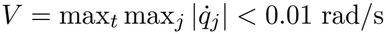 over all 20 joints, with no abort and no invalid step. “Settling” means that *V* and *D* pass. All metrics are computed in double precision at every control sample. The criterion is finite-time and refers to the hand position; *E* is the error of the window-mean position, not the instantaneous error.

#### Initial screening

For each of the three targets and two controller periods, we tested endpoint PD and PID with *K*_*p*_ ∈ {10, 30, 100, 300, 1000} N/m, *τ*_*d*_ ∈ {0.02, 0.1, 0.3} s and *c* ∈ {0, 0.05} . PID also varied *ρ*∈ {0.2, 1, 5} 1/s and the two anti-windup rules. For each endpoint controller (PD or PID), we tested four combinations of gravity compensation (on or off) and posture feedback (on with *K*_*q*_ = 100, off with *K*_*q*_ = 0). This gave eight endpoint controller families in total. The ninth family was the muscle-length reflex, with *k*_*l*_ ∈ {3, 10, 30, 100, 300}, *µ* ∈ {0, 0.01, 0.03, 0.1, 0.3} and *c* ∈ {0, 0.02, 0.05, 0.1}. Together these combinations gave 5640 settings. Each trial lasted 6.2 s (0.2-s transient). We assessed holding over the final 1.2 s.

#### Selection for longer trials

We selected one setting for longer testing from each combination of controller family, target and update period. This gave 54 groups (nine families × three targets × two periods). Within each group, settings that met the holding criterion ranked first. Settings with valid trials that failed the holding criterion ranked next, and settings with invalid trials ranked last. To rank settings within the same category, we divided each holding measure by its threshold and took the largest of the three ratios, max(*E/*0.05, *V/*0.01, *D/*0.005). We preferred the setting with the smaller maximum ratio. If these ratios were equal, we compared the following quantities in sequence: hand position error *E*, mean muscle excitation, the fraction at the upper excitation bound, and the candidate index (the setting ID used in figures and tables). At the first difference, the smaller value ranked higher. Thus every group contributed one setting, including groups with no hold.

#### Long-horizon trials from nine initial conditions

Each selected setting was run for 18.6 s (0.6-s transient) from the nine initial conditions (Sect. 2.2), and the criterion was evaluated in the windows W1 (6.6, 7.8], W2 (12.0, 13.2] and W3 (17.4, 18.6] s (60 samples at 20 ms, 300 at 4 ms). Passing from all nine starts required passing all 27 windows. For the nominal condition, the commands up to 6.2 s were checked online to be identical to the grid rollout.

For settings passing all 27 windows, we also compared each perturbed trajectory with its nominal trajectory at matched times, measuring separation of the fingertips and differences in arm joint angles and velocities (Sect. A.2.2).

#### Physics-step check

To check whether the holding decision depended on numerical integration, we repeated the nominal run for each setting that passed all 27 windows with the physics step halved from 2 to 1 ms, keeping its controller period fixed.

### 2.5 Comparing hand holding in the 34- and 63-muscle models

We asked whether controller settings that held the hand near its target in the 34-muscle model also did so in the 63-muscle model, and how the answer depended on which joints had to remain still.

#### Matching targets and controllers

We used the 63-muscle myoArm model from the same MyoSuite installation, retaining all 38 hinge joints and the same 11 joint equality constraints. Of its 27 independent joints, nine have the same names and ranges as in the 34-muscle model. The other 18 allow thumb metacarpophalangeal and interphalangeal flexion and, in each of the other four fingers, metacarpophalangeal flexion and abduction and proximal (pm) and distal (md) interphalangeal flexion. At each target, we matched the nine shared independent joint angles and set the 18 additional angles to their model default of zero (the lower limit for 12 of them). We used the same constraint polynomials for dependent joint positions and their time derivatives for velocities. The index-fingertip site (IFtip) lies on the distal phalanx and moves with the index finger. Its position at the target posture *q*^*\**^ coincided in the two models at all three targets.

We retained the controller laws, parameter grid, update periods and solver settings and used the same 2-ms physics step for the main comparison. Quantities that depend on the model were computed for its own anatomy. For 63 muscles, we recomputed the inverse-statics excitation *u*_0_; the largest residual joint acceleration at the target postures ranged from 3.8 to 5.3 rad/s^2^. The generalized force *Q* has 38 components, *T* is 38 × 27, and *J* is 3 × 38. The reflex uses the target musculotendon lengths of all 63 muscles, including those that move the fingers. For endpoint control, the optional posture term acts on all 27 independent joints. Without it, the controller specifies no finger angles: the fingertip force demand produces joint torque demands only along the arm and the index-finger joints that move IFtip.

#### Holding criteria

Both models used the same fingertip position error (*E*), drift (*D*), thresholds and evaluation windows. The primary criterion also required the maximum joint speed (*V*) to stay below threshold across all joints (20 in the 34-muscle model and 38 in the 63-muscle model). To compare performance on the same joint set, we also scored each trial using only the 20 shared joints for *V*, retaining the same fingertip measures. This secondary criterion coincides with the primary one in the 34-muscle model. A separate measure used position error and drift at a carpal site (PL-P3 on the capitate) with the same shared-joint speed limit. Both additional measures were specified before the 63-muscle runs and used only for reporting. Selection used the primary criterion throughout.

#### Transfer and model-specific selection

After screening, we compared the models in two ways. First, we transferred the 11 settings that passed the long trials in the 34-muscle model to the 63-muscle model, retaining their gains, update periods and anti-windup rules (Q1). Second, we applied the same ranking procedure independently in each model and compared the selected settings within each of the 54 groups (nine controller families × three targets × two update periods; Q2). For both comparisons, we used nine starts per setting, 18.6-s runs and the same three evaluation windows. One setting was common to both selections in the 63-muscle model, giving 64 distinct settings and 576 distinct runs. We repeated the nominal start of each setting at a 1-ms physics step. We used the saved 34-muscle results, matching screening trials by setting, Q1 trials by setting and start, and Q2 trials by group and start. Before screening, an independent implementation of the stepping loop reproduced 702 reference runs of 32 updates each bit for bit. The model adapter also reproduced the saved 34-muscle results for the low-target PD with gravity compensation, middle-target reflex and high-target PID settings. The comparison concerns transfer between the two models: muscle paths and moment arms, finger mobility and inertia, and the joints receiving posture feedback also differ.

### 2.6 Simulation checks and sensitivity to muscle mechanics

We verified the muscle activation-to-torque map against finite differences and checked start-state integration. At each nominal target we also linearized joint acceleration with respect to joint position and velocity, with activation fixed. We checked the residual acceleration before interpreting these derivatives: at a non-equilibrium posture they measure local mechanical sensitivity (Sect. A.2.3).

We transferred the 11 holding settings without retuning from the native model (R) to variants with a series elastic element in each muscle. At reference force the extension was 2, 4 or 8% of a model reference length (E02, E04, E08); corresponding damped variants (D02, D04, D08) added a 0.01-s damping time constant. These parameters probe sensitivity to series mechanics. All variants and their native control used a 1-ms physics step. We tested all nine starts in R, E04, D04 and D08 and the nominal start in E02, E08 and D02 (429 runs). Failure to find a supported force-balance solution stopped a run and was reported separately from completed non-holds (Sect. A.2.4).

#### Feedback timing and sustained motion

Post-hoc tests examined oscillations under low-target PD control with gravity compensation (setting 85, Table 2), principally in D08. We shortened the controller period from 20 to 4 ms at fixed gains. In a separate comparison we retained 4-ms updates but delayed application of each excitation by 8 ms: the applied command then had the same mean age as under 20-ms updates (10 ms). We compared early changes in oscillation amplitude at two physics steps. We also halved or removed excitation deviations from each muscle’s recent mean and tracked whether trajectories remained close enough for comparison. The multiplier *γ* scaled each muscle’s deviation from its recent mean: 1 retained the ordinary excitation, 0.5 halved the deviation and 0 removed it. For smaller reductions (*γ* = 0.90 and 0.95), a constant-offset control matched the planned mean change for the 10% reduction. We checked realized excitation and cycle-averaged joint and fingertip shifts before attributing changes in motion to modulation. The matched PID controller with conditional anti-windup (setting 631) tested whether integral action changed excitation. Section A.2.5 specifies the controls, stopping criteria, windows and post-hoc design.

**Table 2:**
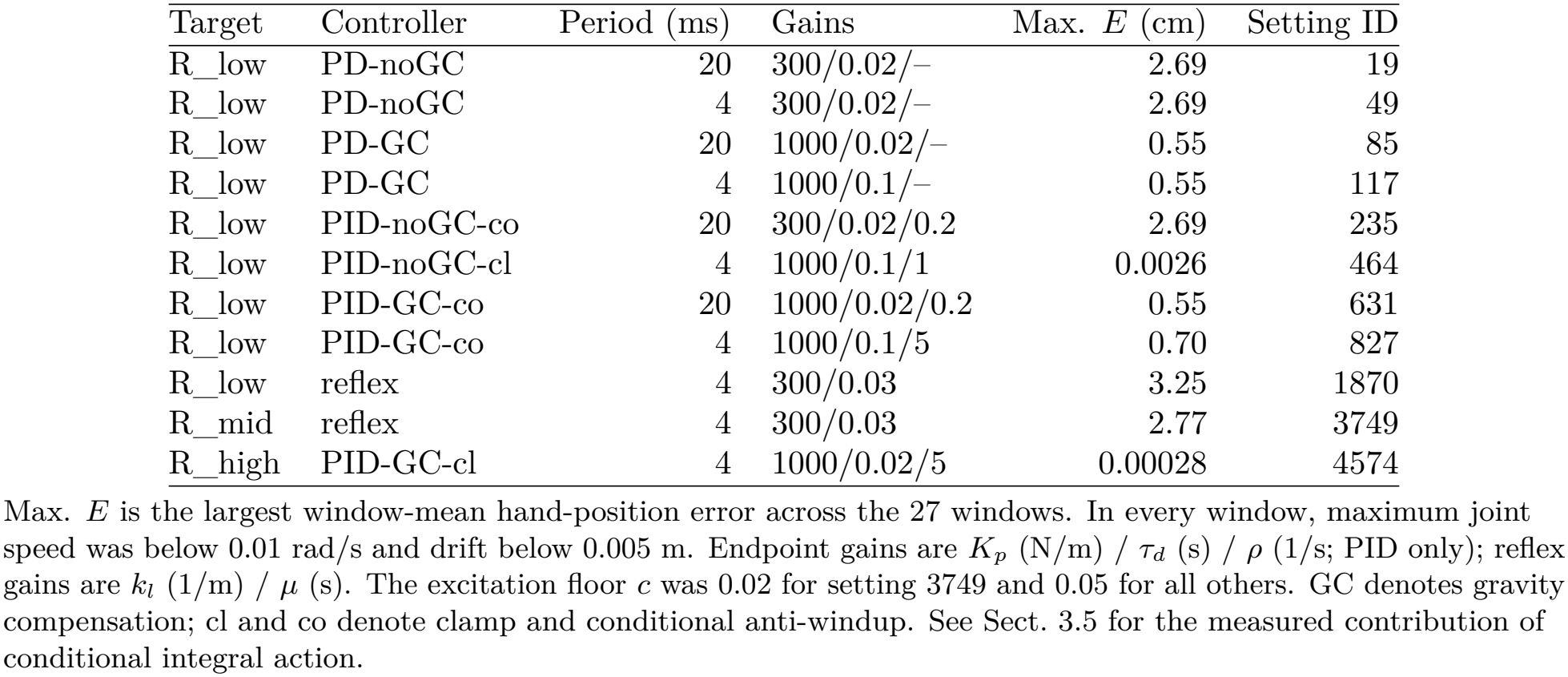
Controller settings that passed the 6.2-s screen and held the hand near the target in all three evaluation windows from each of nine initial conditions.

| Target | Controller | Period (ms) | Gains | Max. *E* (cm) | Setting ID |
|---|---|---|---|---|---|
| R_low | PD-noGC | 20 | 300/0.02/– | 2.69 | 19 |
| R_low | PD-noGC | 4 | 300/0.02/– | 2.69 | 49 |
| R_low | PD-GC | 20 | 1000/0.02/– | 0.55 | 85 |
| R_low | PD-GC | 4 | 1000/0.1/– | 0.55 | 117 |
| R_low | PID-noGC-co | 20 | 300/0.02/0.2 | 2.69 | 235 |
| R_low | PID-noGC-cl | 4 | 1000/0.1/1 | 0.0026 | 464 |
| R_low | PID-GC-co | 20 | 1000/0.02/0.2 | 0.55 | 631 |
| R_low | PID-GC-co | 4 | 1000/0.1/5 | 0.70 | 827 |
| R_low | reflex | 4 | 300/0.03 | 3.25 | 1870 |
| R_mid | reflex | 4 | 300/0.03 | 2.77 | 3749 |
| R_high | PID-GC-cl | 4 | 1000/0.02/5 | 0.00028 | 4574 |

Max. *E* is the largest window-mean hand-position error across the 27 windows. In every window, maximum joint speed was below 0.01 rad/s and drift below 0.005 m. Endpoint gains are *K*<sub>*p*</sub> (N/m) / *τ*<sub>*d*</sub> (s) / *ρ* (1/s; PID only); reflex gains are *k*<sub>*l*</sub> (1/m) / *µ* (s). The excitation floor *c* was 0.02 for setting 3749 and 0.05 for all others. GC denotes gravity compensation; cl and co denote clamp and conditional anti-windup. See Sect. 3.5 for the measured contribution of conditional integral action.

### 2.7 Closed-loop perturbations and constant-excitation mechanics

The closed-loop map *Y*_*k*+1_ = Φ(*Y*_*k*_) advances the arm and controller together over one control period. Its state includes joint positions and velocities, muscle activations and solver warmstart values, plus integral or series-extension states where applicable. We tested candidate fixed points and checked whether perturbations remained on the same numerical and controller branches as their widths decreased. We assessed native and double-precision reference maps separately (Sect. A.2.6).

In a post-hoc test of four rigid holding settings (low- and middle-target reflex, low-target PD and high-target PID), we used one-coordinate perturbations to estimate a linear prediction 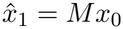, where *x*_*k*_ is the scaled full-state difference from an unperturbed run. We used paired perturbations (*M*_two_) or, at activation bounds, the feasible one-sided response (*M*_bound_). At each of four solver settings (L0–L3, with successively tighter convergence tolerances), we tested agreement across perturbation amplitudes of *r* = 16, 8 and 4 times each coordinate’s base size, prediction in 16 independent mixed directions and prediction over 0.2 and 2 s. These tests address finite-amplitude prediction. In a prospective test in the PD and PID settings, leading right singular vectors selected initial perturbations for one update, 0.2 s and 2 s. At the primary amplitude (*r* = 8) the measured gain had to exceed or equal 1.25 times the maximum in each of two comparison groups (fixed patterns and a pseudorandom direction; Sects. A.2.7 and A.2.11).

After the pilot stiffness measurement (C0), we used smaller difference increments and a wider range of probe forces (C-R). At one verified equilibrium per target, we estimated static horizontal hand stiffness 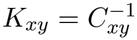 from the displacement per unit applied load with activation fixed and vertical motion free. We checked the estimate against independent equilibrium displacements and known spring additions. In a separate post-hoc test (C0-S) at the middle- and high-target equilibria, we removed feedback, held excitation at the equilibrium activation *a*_*h*_ and perturbed joint position and velocity along the leading mechanical eigenvector. We compared both perturbation signs with the unperturbed trajectory over 0.5 s at physics steps of 1 and 0.5 ms. Activation perturbations were not tested. This tests the mechanics under constant excitation; closed-loop predictions use Φ above (Sect. A.2.12).

## 3 Results

In the 34-muscle model, 163 of 5640 settings passed the initial screen. Eleven selected settings subsequently held across all 27 windows of the nine-start test. When transferred to the 63-muscle model, these settings held in 89 of 99 runs when joint speed was assessed over the shared joints, and in 18 of 99 when all joints had to remain still (Sect. 3.7).

### 3.1 Implementation verification and target sensitivity

Starting from the same prescribed state, the environment and an independent integration loop agreed bit for bit in joint position, velocity and muscle activation in six tests (one and five control steps at each of the three targets). The largest relative Frobenius difference between the analytic and finite-difference force maps in calibration was 3.6 × 10^−15^. In static probes, the reduced-coordinate allocation reproduced the direction and magnitude of the demanded generalized force (cosine *≥* 0.99999999, magnitude ratio 0.99999–1.000001).

At the targets under the approximate inverse-statics command, the joint acceleration is not zero (largest 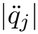 over the 20 joints 2.5–3.4 rad/s^2^), so these points are not equilibria. The mechanical linearization in Sect. 2.6 has dimension 40 × 40. Its largest real part is 1.0–2.8 1/s over the three targets and changed by less than 0.002 1/s between the two finite-difference step sizes. We report this only as a local sensitivity of a non-equilibrium state; it is not a linearization in the full state (*q*, 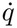, *a*), and it is not evidence about the stability of an equilibrium or of the closed-loop holds below. These non-equilibrium target diagnostics are distinct from the constant-excitation equilibrium test in Sect. 3.6.

### 3.2 Hand holding at the three targets

At every target at least one fixed-parameter controller held the hand (Fig. 2): a PD controller with gravity compensation at R_low (20 ms), the reflex at R_mid (4 ms) and a PID controller with gravity compensation and clamp anti-windup at R_high (4 ms). In the last window of the 18.6-s nominal runs, *E* was 0.55, 2.77 and 1.6 × 10^−7^ cm; maximum joint speed in all three examples was well below the holding threshold of 0.01 rad/s. The PD and reflex examples thus retain offsets within the prespecified 5-cm tolerance, whereas the clamp-PID example has a near-zero offset; passing this criterion does not by itself imply precise target tracking. The joint configuration, however, did not return to *q*^*\**^: the largest mean deviations of an arm joint were − 20.7° (flexion), − 44.3° (pronation–supination) and − 16.9° (flexion). Of the 18 settings selected per target for the long-horizon trials, nine passed all 27 evaluation windows at the low target, compared with one at the middle target and one at the high target (Table 2). Across these 11 settings, the largest mean arm-joint deviation ranged from 3.2° (reflex, R_low) to 50.6°. These are therefore examples of holding the hand, not of holding the joint posture.

**Figure 2:**
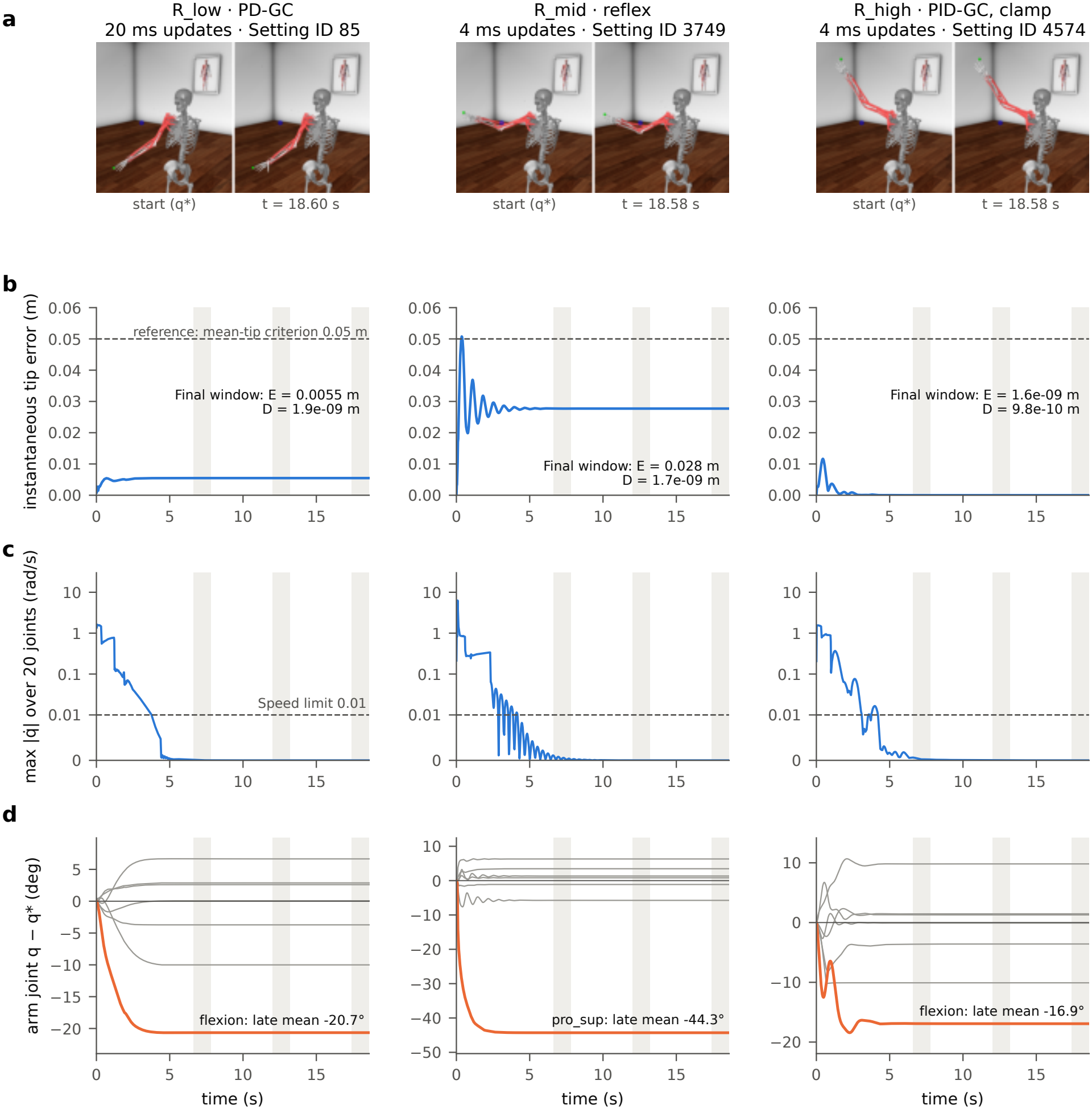
Nominal-start holds at three targets. GC denotes gravity compensation at the nominal target posture. **a** Start and last saved configuration. **b** Instantaneous hand error with the windows (gray); the dashed line is the 0.05-m reference level of the window-mean criterion, not a pass condition for the instantaneous error. **c** Maximum joint speed. The axis is linear up to the holding threshold (dashed, 0.01 rad/s) and logarithmic above it. **d** Arm-joint deviation *q* − *q*^*\**^ (no wrapping); the joint with the largest late-window mean deviation is highlighted. Traces are saved every 0.02 s; decisions use full-rate double precision values.

### 3.3 Finite grid and paired comparisons

Of the 5640 grid settings, 163 met the criterion in the 6.2-s runs, in 11 of the 54 groups (Fig. 3a): all PD/PID endpoint groups at R_low, the reflex at R_low and R_mid at 4 ms, and the PID-GC endpoint group at R_high at 4 ms. No setting with the posture term held. These are counts on a finite grid that deliberately includes poor gains, not probabilities of success.

**Figure 3:**
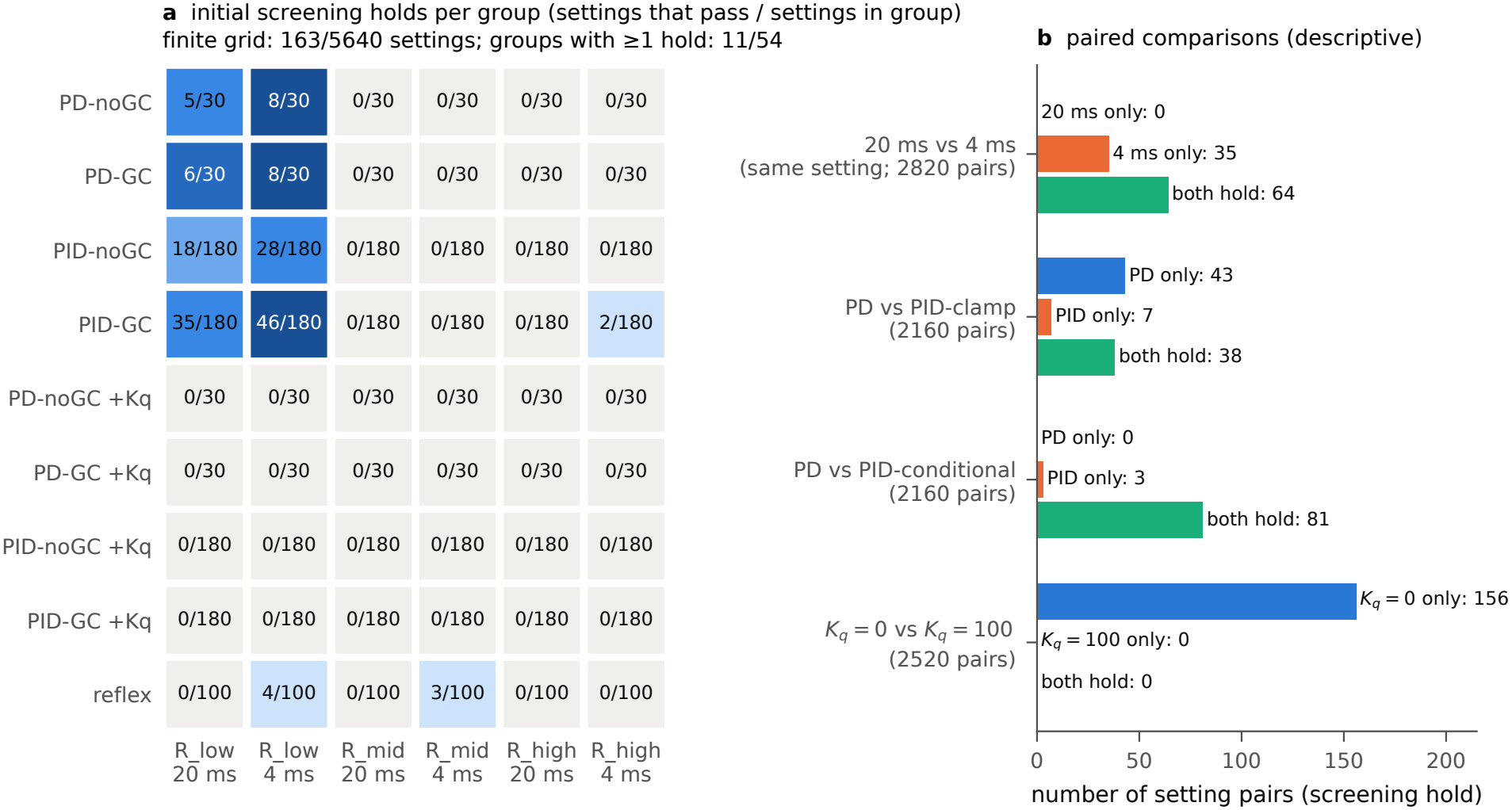
Hand holding in the 34-muscle model. A setting passed if it met the hand-position, drift and joint-speed criteria in the final 1.2 s of the 6.2-s trial. **a** Columns pair each target posture (low, middle or high) with a controller update period (20 or 4 ms). Cells show settings passed/tested; darker shading indicates a larger fraction passing. GC and noGC denote endpoint control with and without gravity compensation at the target posture. Rows marked +*K*_*q*_ also include joint-posture feedback (*K*_*q*_ = 100 N m/rad). **b** Bars count matched pairs in which only the first setting passed, only the second passed, or both passed. Pairs in which neither passed are omitted. Clamp and conditional denote the two PID anti-windup rules. Each PD setting was paired with three PID integral gains per rule, so the pair counts are descriptive.

Paired comparisons of settings that differ in one factor (Fig. 3b) were asymmetric in this grid: 35 settings held only at 4 ms and none only at 20 ms (64 held at both); PD held where the matched clamp-PID did not in 43 pairs and the reverse occurred in 7 (38 both); for conditional PID the counts were 0 and 3 (81 both), and in 455 of 2160 pairs the conditional-PID command sequence was identical to that of the matched PD. Adding the posture term removed all 156 holds of the matched *K*_*q*_ = 0 settings. Among pairs in which both settings held, gravity compensation reduced *E* in all 13 matched PD pairs at R_low (same *K*_*p*_, *τ*_*d*_, *c* and Δ*t*; ratio of *E* with to *E* without gravity compensation 0.48–0.66), and clamp PID reduced *E* relative to the matched PD in all 38 pairs (ratio 5.6 × 10^−4^ to 0.67). At R_high, both settings that held in the 6.2-s screen used PID with gravity compensation, clamp anti-windup and a 4-ms update period. For the setting selected for the long-horizon trials, the matched PD did not settle in the grid run (*V* = 0.048 rad/s). Because each PD setting is the reference for several pairs, these counts are descriptive and are not tested statistically.

### 3.4 Persistence and initial conditions

The 11 settings selected from the groups that contained a hold are exactly the 11 settings that passed all 27 windows of the long-horizon check (Fig. 4, Table 2). None of the 486 long-horizon rollouts was invalid or unsupported. Of the 43 settings selected from groups without a 6.2-s hold, three passed the late window at 18.6 s: the high-target PD and PID settings without gravity compensation (IDs 3817 and 4223) failed only W1 at every start (18/27 windows passed). The low-target reflex setting (ID 1771) failed only W1 after the positive shoulder-elevation angle perturbation (26/27 passed). In each failed window, only joint speed exceeded its threshold. The 152 unselected settings that held for 6.2 s were not run for 18.6 s. In the nominal condition, two PID settings that held from all nine starts (235 and 631) produced command sequences identical to those of PD settings (19 and 85, respectively), so the 11 holding settings comprise nine distinct nominal command sequences. This equivalence depends on the initial condition: across the nine conditions the 11 settings produced 8 to 11 distinct sequences.

**Figure 4:**
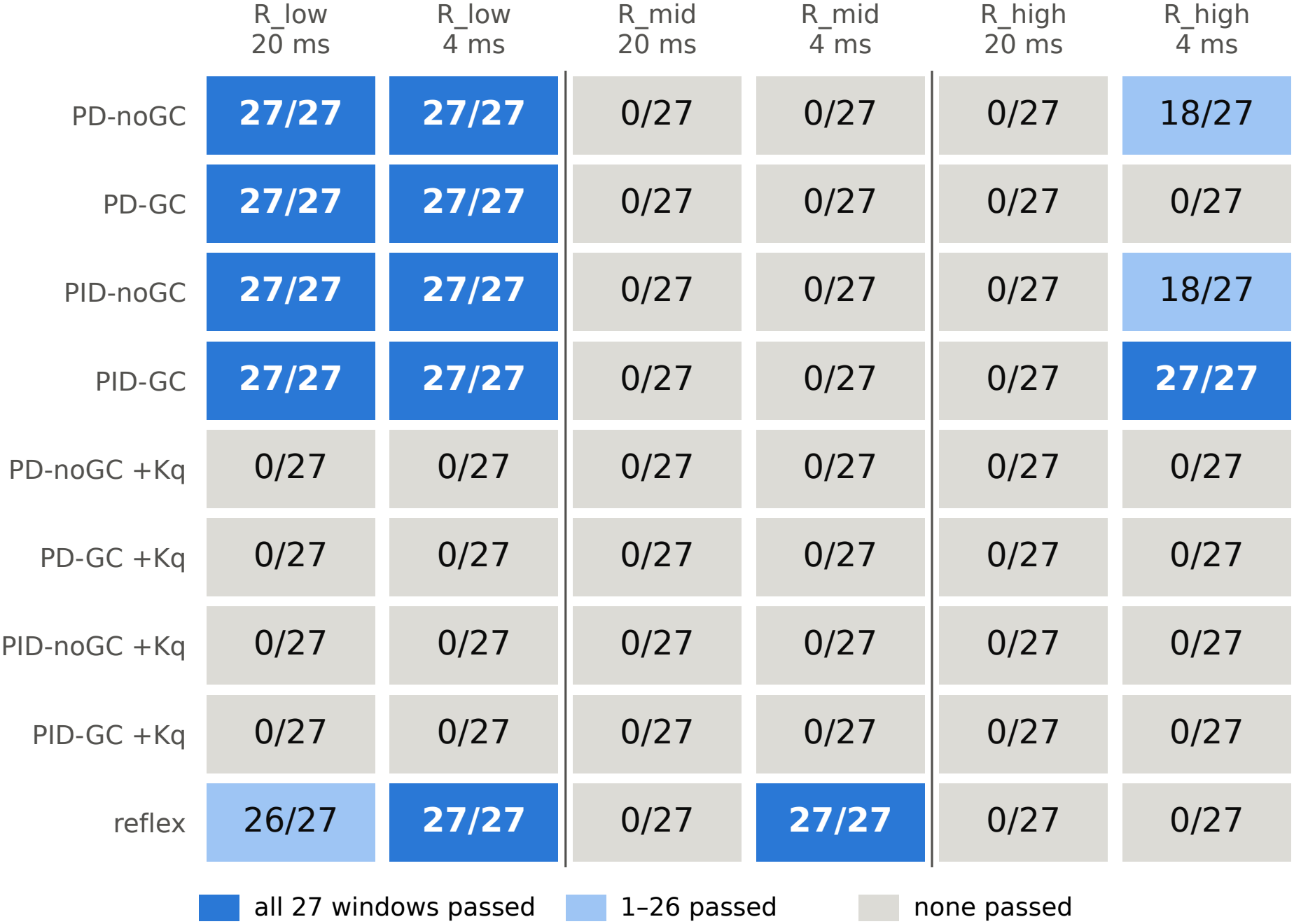
Long-horizon holding across initial conditions. Each cell corresponds to the same group as in Fig. 3a, where the fraction counts settings that passed the short trial. Here one setting per group was selected by the prespecified rule. Each fraction shows windows passed out of 27 for that setting (three windows in each of nine 18.6-s runs from different initial conditions). Windows do not cover the intervals between them, and passing the tested conditions does not establish stability under arbitrary perturbations.

To see whether the effect of a small change in the initial state diminished during holding, we compared each perturbed trial with the nominal trial at the same elapsed time, keeping the controller setting fixed (11 settings, 88 pairs; Fig. S1). In every pair, the W3 maximum of each of the hand-position, largest arm-joint angle and largest arm-joint speed differences was below its value at the first stored sample, one controller period after initialization. Some runs retained late-window differences: the largest W3 mean residuals across pairs were 2.7 mm and 0.27° (both in setting 827). The largest sampled hand separation was 19.0 mm across all perturbations and 5.3 mm for velocity perturbations only. Following the first sample, hand separation increased in 10/44 position and 40/44 velocity perturbations. These are finite-time differences of thinned single-precision traces; they do not establish convergence to a common state or closed-loop asymptotic stability.

With the physics step halved, all 11 holding settings again passed W1–W3 in the nominal condition. Over W1–W3 the largest changes were |Δ*E*| = 5.8 × 10^−6^ m, |Δ*V*| = 1.6 × 10^−4^ rad/s and |Δ*D*| = 1.5 × 10^−5^ m (the latter two in W1; in W3, 5.0 × 10^−7^ rad/s and 1.2 × 10^−8^ m). This is a two-level sensitivity check of the nominal runs only; the perturbed conditions and the rest of the grid were not rerun at *h* = 1 ms.

### 3.5 Remaining failures and inactive integral action

Figure 5 illustrates two kinds of failure among the selected settings. First, in 16 of the 54 selected settings (nominal condition), mostly endpoint controllers at R_mid and R_high, the arm settled (*V ≤* 1.1 × 10^−5^ rad/s) but the hand stopped 5.3 to 30 cm from the target, with *E* almost the same in W1, W2 and W3. Second, all settings with the posture term (*K*_*q*_ = 100) kept moving (late-window *V* 0.47–6.6 rad/s in the nominal runs). At 4 ms the reduced demand on the seven arm joints was approximately realized (late-window mean relative residual 1.1 × 10^−5^ to 0.029 over the 12 posture-term settings at 4 ms; 1.7 × 10^−5^ in the example of Fig. 5b1), whereas the demand on the two thumb carpometacarpal joints, which were at their soft range limits, was not realized; at 20 ms the arm-joint residual was 0.56–0.72. The cause of the sustained arm motion was not identified.

**Figure 5:**
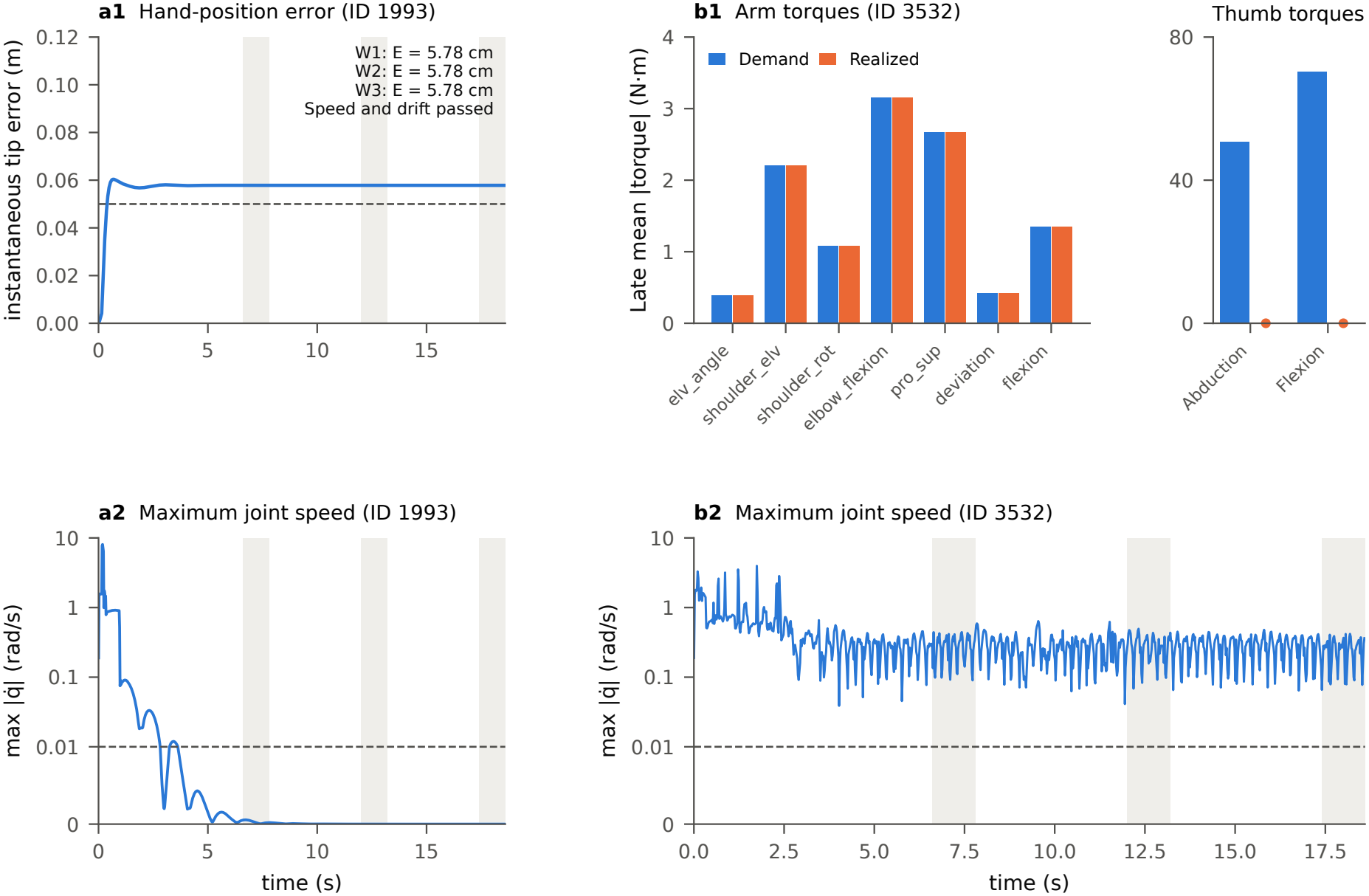
Two holding failures at the middle target from the nominal start (4-ms updates). Left: with PD-GC (setting 1993), the hand remained outside the target tolerance (**a1**) even after joint speed fell below threshold (**a2**). Right: with PID-GC and posture feedback (*K*_*q*_ = 100, setting 3532), late-window realized torques matched demand at the arm joints but fell far short at the thumb (**b1**), and joint speed remained above threshold (**b2**). The lower graphs show the largest absolute speed across all 20 joints at each saved time, on a common axis that is linear up to the holding threshold (dashed, 0.01 rad/s) and logarithmic above it. Arm and thumb torque axes span 0–4 and 0–80 N m, respectively; dots mark near-zero realized thumb torques.

In the 10 selected endpoint settings with conditional anti-windup the integrator was frozen in almost every control step (frozen fraction 0.994–1.000, median 0.999, over their 90 long runs). Three of these settings (235, 631, 827) held from all nine starts and seven did not, so the freezing limits the integral action of this rule but does not by itself determine whether a setting holds. In the nominal low-target trial, conditional PID (setting 235) and its matched PD (setting 19) both held and sent identical muscle excitations throughout the trial.

### 3.6 Plant sensitivity and local diagnostics

#### Transfer to series-elastic muscle models

Of 429 scheduled 1-ms cases, 328 completed 18.6 s and 305 held in all three windows (Table 3). The 23 completed non-holds comprise 22 D08 cases and one E08 case. The other 101 stopped when the solver could no longer follow a continuous force-balance solution for the series element. R and D04 each held in all 99 cases. E04 completed and held in 14 of 99, with 85 such early stops; D08 completed all 99 and held in 77. Coverage differs between the nominal-only and nine-start variants, so these counts are not success-rate estimates over a common initial-condition distribution. The six predefined half-step cases include five completed holds and one domain stop, separately from the 429 main cases.

**Table 3:** Holding outcomes by muscle model with controller parameters fixed (18.6-s trials, *h* = 1 ms). R is the native model; E and D have series compliance without and with serial damping. At the reference elastic force, the series element extends by 2%, 4% or 8% of the reference length in models ending in 02, 04 or 08, respectively. Each trial tested one controller setting from one initial state. All models used the same 11 settings, with nine starts per setting for R, E04, D04 and D08 and only the nominal start for E02, E08 and D02. Early stops occurred when the solver could no longer follow a continuous solution for force balance in the series element. The six predefined half-step checks are separate.

|  |  | Completed trials |  |  |
|---|---|---|---|---|
| Model | Total trials | All 3 windows passed | At least 1 window failed | Stopped early |
| R | 99 | 99 | 0 | 0 |
| E02 | 11 | 5 | 0 | 6 |
| E04 | 99 | 14 | 0 | 85 |
| E08 | 11 | 0 | 1 | 10 |
| D02 | 11 | 11 | 0 | 0 |
| D04 | 99 | 99 | 0 | 0 |
| D08 | 99 | 77 | 22 | 0 |
| Total | 429 | 305 | 23 | 101 |

In D08, the two low-target controllers described in Sect. 2.6 showed persistent motion near 7 Hz at all nine starts: the PD controller with gravity compensation (setting 85) and its PID counterpart with conditional anti-windup (setting 631). At four starts of the high-target PID setting, joint speed exceeded the threshold only in W1 (Fig. S2b). The one completed E08 non-hold also fails W1 rather than all late windows. These observations establish neither a limit cycle nor the cause of the motion. Only the PD controller underwent the half-step check.

#### Changing the controller update period

From the nominal start in D08, shortening the PD update period from 20 to 4 ms restored holding in all three windows at both physics steps. Wrist and arm speed fluctuations fell to less than 0.2% of their respective baseline amplitudes. The rigid model and D04 already held at both update periods (Fig. S2, Table A1). In a separate low-target comparison in the rigid model, PD setting 87 (20 ms) failed all three windows, whereas its matched setting 117 (4 ms) passed all three at both physics steps.

Adding an 8-ms delay to the 4-ms updates slowed the early decay of D08 oscillations. Strong reductions in excitation modulation moved the arm away from the posture used for comparison. A smaller reduction lowered oscillation amplitude and frequency, but also shifted joint posture, leaving the mechanism unresolved. At 4 ms the conditional PID controller reproduced the PD trajectory with zero integral contribution (Sect. A.2.13).

#### Can local feedback dynamics predict the motion?

The fixed-point tests yielded no validated local stability spectrum for the closed-loop system. In four rigid-model settings, maps fitted to finite perturbations also failed to predict the full response to independent perturbations (Table A2). In two settings the maps nevertheless identified directions that produced relatively large responses: seven of nine exceeded both comparison groups by the prespecified 25% margin (Sect. A.2.13).

#### Directional stiffness of the fingertip

With activation fixed, we obtained a verified stiffness measurement at one equilibrium per target. In the horizontal measurement plane the symmetric stiffness matrix was positive definite at the low target; at the middle and high targets it had one negative eigenvalue (Table A3). Eight additional starts at the middle target led back to the same equilibrium, leaving no second equilibrium there at which to compare stiffness.

#### Mechanical response after feedback removal

At the middle and high targets we removed feedback and held muscle excitation at its equilibrium value. Small joint displacements along the leading mechanical mode then grew by factors of approximately 7.71 and 1.84 over 0.5 s, respectively, in agreement with the linear predictions for both perturbation signs and physics steps (Fig. 6). These tests concern the response of the mechanics at constant activation; the same settings held the hand near the target with feedback enabled.

**Figure 6:**
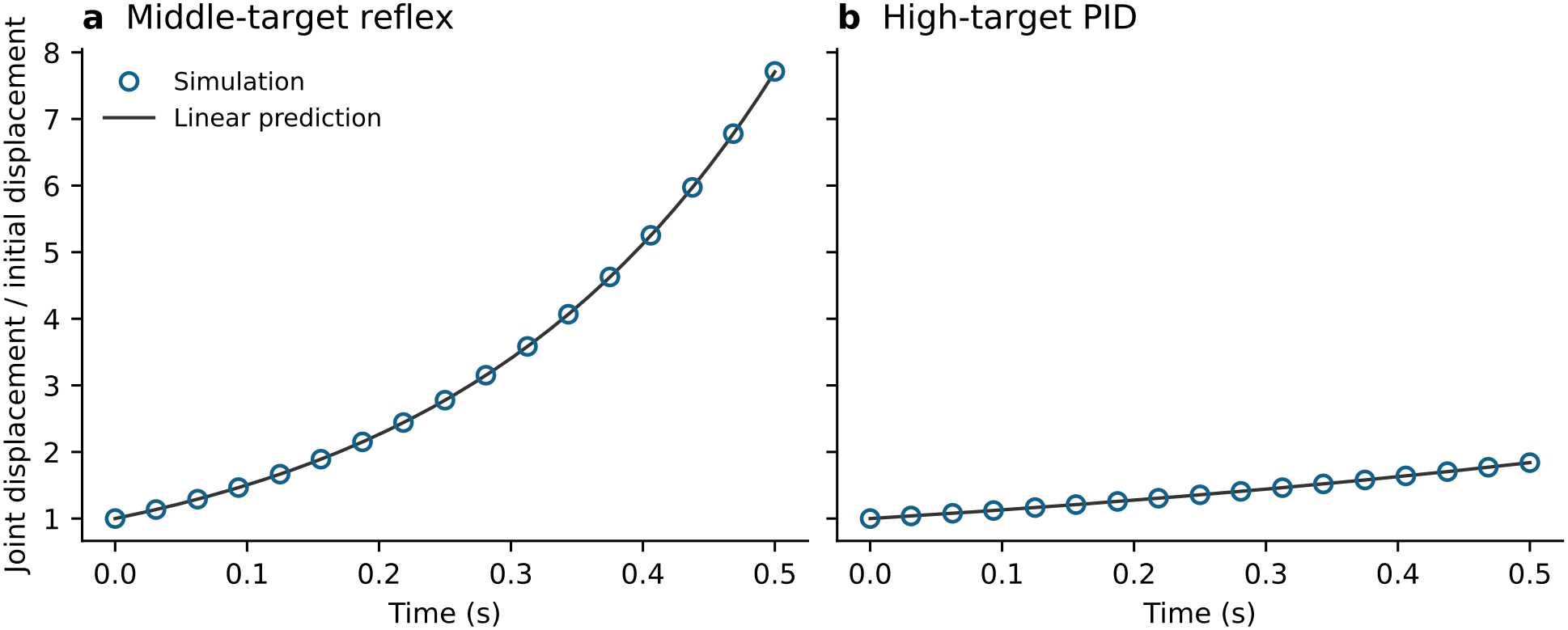
Growth of joint displacement after perturbation at two equilibria with muscle excitation held constant and feedback removed. **a** Middle-target reflex setting. **b** High-target PID setting. Each curve shows the norm of the joint displacement from the matched unperturbed trajectory divided by its initial value. Both perturbation signs along the leading mode and both physics steps were tested; the largest initial joint-angle change was 10^−6^ rad. Open circles show selected simulation samples, alternating among the four runs at their recorded times. The solid line shows the linear prediction exp(*αt*). Across all samples, the maximum relative difference from the prediction was below 0.08% at the middle target and below 0.003% at the high target.

### 3.7 63-muscle comparison

When the 11 selected settings were transferred without retuning (Q1), 89 of 99 trials met the same fingertip and shared-joint criteria as in the 34-muscle model across all three windows (the prespecified secondary criterion). Under the prespecified primary criterion, which assessed speed at all 38 joints, 18 of the same 99 trials passed (Table 4). Nine settings passed the shared-joint criterion from all nine starts, one from eight and one from none (Table A4).

**Table 4:** Hand holding in the 34- and 63-muscle models. Each entry gives trials passed/total. All columns use the same fingertip error and drift criteria. The joint-speed criterion applies to all joints (primary) or, in the 63-muscle model, to the 20 shared joints (secondary).

|  | 34-muscle | 63-muscle |  |
|---|---|---|---|
| Trials | All 20 joints | All 38 joints | Shared 20 joints |
| Initial screen (6.2 s) | 163/5640 | 4/5640 | 104/5640 |
| Fixed transfer (Q1): 11 settings from the 34-muscle model | 99/99 | 18/99 | 89/99 |
| Model-specific selection (Q2): 54 settings per model | 107/486 | 18/486 | 27/486 |

In Q1 and Q2, each setting was tested from nine starts for 18.6 s; a trial passed if all three windows passed. The two sets of trials overlap and are reported separately. Table A4 gives the Q1 results by setting.

Under the primary criterion, 4 of the 5640 settings held in the 63-muscle grid (Table 4), compared with 163 in the 34-muscle grid: reflex settings 1870, 1874, 3749 and 3750 at the 4-ms period, two at R_low and two at R_mid. Their *E* was 0.037–0.045 m, 5–13 mm inside the tolerance. No R_high setting and no setting with the posture term held. One run (setting 1399) stopped after a simulator reset and is counted as a non-hold. Under the secondary criterion 104 settings held (56 at R_low, 10 at R_mid and 38 at R_high), again none with the posture term. Of the valid settings, 622 met the *E* and *D* criteria but failed the 38-joint speed criterion; 100 of these met the shared-joint speed criterion, so the secondary count consists of the 4 primary holds and these 100. All 100 were endpoint settings; for the reflex the two criteria gave the same four settings. The separate carpal-site measure gave 46. Of the 163 34-muscle holds, the same setting also held in the 63-muscle grid in 3 cases under the primary and 51 under the secondary criterion; 53 of the 104 secondary holds were settings that had not held in the 34-muscle grid (38 at R_high). The secondary count is therefore not the 34-muscle count with finger effects removed.

Selection with the unchanged primary key picked different settings in 51 of the 54 groups. In Q2, 18 of the 486 runs passed all three windows under the primary criterion (settings 1874 and 3749 from all nine starts) and 27 under the secondary criterion (adding R_high PD-GC setting 3841), compared with 107 runs in the 34-muscle model, including all nine starts of its 11 holding settings. Settings were not reselected with the secondary key.

In each of the 71 fixed-transfer (Q1) runs that failed only the primary criterion, the failure was confined to W1 and to joint speed. The fastest joint was the middle-finger distal interphalangeal joint md3 (0.22–0.32 rad/s); the window-mean hand error was 0.003–3.6 cm (median 1.2 cm), *D* was at most 0.065 mm, and W2 and W3 passed the primary criterion. The 18 added finger coordinates started at zero, and these endpoint settings specified no finger-angle targets. In all 71 saved trajectories the distal middle-finger angle first came within 0.01 rad of its upper flexion limit between 7.18 and 11.48 s. It remained within that distance in every saved sample from 12 s onward (examples in Fig. S3). The other failures were not confined to the fingers: setting 827 from the elbow-flexion *q*− start moved its shared joints at 0.94–1.06 rad/s in all windows (fastest: shoulder rotation), and in setting 4574 the hand error was below 0.02 mm while the shared joints moved at 0.033–0.046 rad/s. The primary long-horizon holds comprise 27 distinct runs (settings 1870, 1874 and 3749 from all nine starts), because the 18 Q1 and 18 Q2 passes share the nine runs of the shared middle-target reflex setting. Grid-holding setting 3750 was not selected and was not run for 18.6 s. At *h* = 1 ms the nominal runs of these three settings again passed all windows; the perturbed starts were not rerun at that step.

Figure S3 illustrates the fingertip and finger trajectories for three nominal starts. As in the 34-muscle grid, conditional PID often produced the same command sequence as its matched PD (in 426 of 2160 pairs in the 63-muscle and 455 in the 34-muscle model).

## 4 Discussion

### What the results show

Fixed-parameter feedback held the hand near all three targets, with gains selected for each target and update period. Of the 99 transfer trials, 89 met the prespecified secondary criterion for fingertip holding and speed at the 20 shared joints, and 18 of the same 99 met the prespecified primary criterion requiring all 38 joints to stay below the speed limit (Sect. 3.7).

### What the controllers use

“Fixed-parameter” refers to time-invariant gains. The PD and PID controllers use an exact, state-dependent static force map of the plant at every control step, and the reflex uses the target musculotendon lengths. The successes therefore rely on accurate static information about the plant; they say nothing about whether such information could be learned, or about how the nervous system might obtain it. In human reaching experiments, participants adapted arm impedance to a destabilizing force field [22]. We make no claim that learned or predictive control is required, nor that a static internal model is unnecessary.

### Plant and controller dependence

Because conclusions depend on the periphery model [2], the native results are supplemented by the frozen-controller series-model tests above. D04 preserved the tested holds, whereas D08 added sustained motion and the undamped variants encountered force-equilibrium domain limits. These limits and the mixed initialization are part of the declared sensitivity model, not controller instabilities or evidence that anatomical compliant tendons cannot support holding. The tests neither identify optimal gains for each plant nor isolate the cause of the approximately 7-Hz motion. Günther et al. found that a load attached to an undamped series elastic element oscillated at approximately 11–19 Hz; adding a damper in parallel with that element, with damping increasing with muscle force, suppressed the vibrations [3]. Our D08 model used damping proportional to series stiffness and exhibited motion near 7 Hz under feedback. These tests do not identify the load-oscillation mechanism in D08. Halving the native physics step addresses numerical integration, not anatomical model validity. Other models and controllers have produced stable movement and posture [4, 5, 6, 7], and the present results agree with them in that fixed feedback can hold. They differ from the present set-up in ways that may matter for holding. All four used compliant tendons, three with a serial damping element [7, 6, 5]. Their reflexes were combined with feedforward stimulation and, in two of them, with feedback delays of 25 ms or more [4, 5]; Walter et al. set sensor delays to zero [7]. Gains were optimized to trajectories or data [4, 5], or chosen heuristically or fixed [7, 6]. The tasks were movements, stance or spinal posture; where a posture was held after a movement, the hold lasted up to 2 s [6, 5]. Kistemaker et al. also reported marked endpoint oscillations with reflex-only control at high gains [4]. The native model has inelastic paths without serial damping; our controllers have no feedback delay other than the sample-and-hold of the controller period, and our gains were selected per target from a grid; and holding was judged over 18.6 s. These differences are confounded, so the comparison does not isolate the effect of any one of them. The D08 interventions tested whether command timing and excitation modulation affected the sustained motion. At unchanged gains, more frequent updates restored holding at the nominal start. Delaying command delivery while keeping the faster updates moved the early rate of amplitude change toward that of the slower-update control, although the later motion differed. Command timing thus affected the early response. Reducing excitation fluctuations by 10% reduced oscillation amplitude and frequency at all four sampled phases. Matching the mean excitation change alone produced little amplitude change, but the two interventions also shifted joint posture differently. The data therefore leave open whether the reduction acted directly through feedback or through the change in posture. The constant-excitation tests at two equilibria of the rigid model address a separate question and do not establish the cause of the motion in D08.

The selected settings kept the index fingertip near its target in most transfer trials even though the index finger could now bend. Middle-finger flexion, by contrast, does not itself displace the index fingertip. In the 71 trials that passed only the shared-joint criterion, the endpoint controllers specified no finger-angle targets: the index fingertip met the holding criterion while the middle finger was still settling in the first window. All joints met the speed limit in both later windows. For the reflex, which also acts on finger muscles, the same settings passed both criteria. The tested posture term, which does set targets for all independent joints including the fingers, produced no hold under either criterion, so an explicit finger target is not shown to be sufficient. A dedicated redesign to hold the hand and finger posture together was not tested and is left for future work. Because the models differ in muscles, moment arms, finger joints and their inertia, and posture-term coordinates at once, the comparison does not attribute differences to muscle number.

### Implications for controller design

First, specify which joints must remain still alongside the hand-position objective. The hand could remain near its target even when joint angles differed across starts or a finger was still moving (Sects. 3.4 and 3.7).

Second, compare requested torques with those the bounded muscle excitations actually deliver. The thumb-joint demands were poorly realized in the posture-feedback trials. Even where the arm-joint demand was approximately realized, motion could persist (Fig. 5b2).

Third, examine the timing and the contribution of each feedback term. Conditional anti-windup often suppressed integral action, so naming a controller PID does not establish that its integral term affected the trajectory (Sect. 3.5). In D08, shortening the update period restored holding at unchanged gains from the nominal start (Sect. 3.6).

The full-state follow-up places a further limit on local linear explanations. Across the tested amplitudes, the fitted maps failed to predict the full response to independent perturbations. A near-unit spectral radius of such a map cannot supply a measured decay time. Yet its leading singular directions could still identify perturbations with relatively large measured responses against the specified comparisons. Selecting a direction and predicting the subsequent trajectory are distinct uses of a local model; success at the former does not validate the latter.

### 4.1 Limitations

The main controller comparison uses one rigid-tendon 34-muscle model, per-target gains, finite-time windows and small single-joint offsets; holding refers to the hand. The main comparisons used sampled feedback without a physiological conduction delay. PD/PID use a model-based static force map, not model-free control. The added series variants are sensitivities, not validated anatomical models; strict domain stops, mixed initialization and unequal start coverage are disclosed.

Finite-amplitude predictions were tested in four rigid settings; the prospective direction test covered two settings at one solver level with small comparison groups. An infinitesimal closed-loop spectrum and asymptotic stability remain unverified. The constant-excitation test covered two rigid equilibria; the wrist-limit case and the elastic-model mechanical spectra remain unresolved.

The tested starts yielded only one distinct equilibrium per target for stiffness measurement. Even the larger, hand-position-preserving perturbations at the middle target returned to the same base (Sect. 3.6). Stiffness sensitivity across distinct bases at the same target, elastic and closed-loop stiffness, and anatomical or human stiffness validation remain untested. A positive xy stiffness in one setting is not proof of full-state stability; indefinite matrices are not plotted as ellipses. The 63-muscle comparison changes several plant properties together and does not separate the contributions of muscles, moment arms, finger joints or inertia. Its long-horizon tests cover only the selected and transferred settings (grid-holding setting 3750 was not tested), its half-step check covers nominal starts only, and the secondary criterion was not used to reselect settings. The full-state, stiffness, constant-excitation and series-model analyses were performed only in the 34-muscle model. The D08 interventions were post hoc. The update-period test used the nominal start, the delay comparison used one saved state per physics step, and the weak-modulation test sampled four phases of one trajectory. Strong reductions left the comparison domain before the evaluation window. These experiments constrain explanations for motion along the tested trajectory; they do not identify the mechanism that sustains it.

## 5 Conclusion

Fixed-parameter feedback could hold the hand near each of the three targets in the 34-muscle arm model. Eleven target-specific settings met the holding criterion in every evaluation window of 18.6-s trials from all nine tested initial conditions.

The transferred settings maintained near-target holding in 89 of 99 trials under the shared fingertip and joint-speed criteria. Under the prespecified primary criterion, which also required the additional joints to remain still, 18 of the 99 trials passed all three windows.

In two successful settings of the 34-muscle model, small perturbations along a mechanical mode grew over 0.5 s when excitation was held at its equilibrium value and feedback was removed. Thus finite-time holding under feedback can coexist with a mechanically expanding direction under constant excitation. The local stability of the full closed-loop system remains unresolved.

## Declarations

### Use of AI-assisted tools

The author used OpenAI Codex and Claude Code to assist with programming, analysis checks, manuscript drafting and editing.

### Data and code availability

The analysis code and saved results are available in GitHub release v0.2.0. The data attachment includes the original and revised classifications of the elastic-model perturbations. The source archive includes an index of data files and their checksums. Zenodo archives both the software (doi:10.5281/zenodo.24457722) and the dataset (doi:10.5281/zenodo.24458113) for this release. Software versions were NumPy 2.4.6, SciPy 1.17.1, MuJoCo 3.6.0, MyoSuite 2.12.2 and Gymnasium 1.2.3.

## A Supplementary figures, methods and tables

### A.1 Supplementary figures

Figures S1 and S2 show the time course of responses to initial perturbations and to changes in controller timing. Figure S3 shows finger motion during transfer to the 63-muscle model.

**Figure S1:**
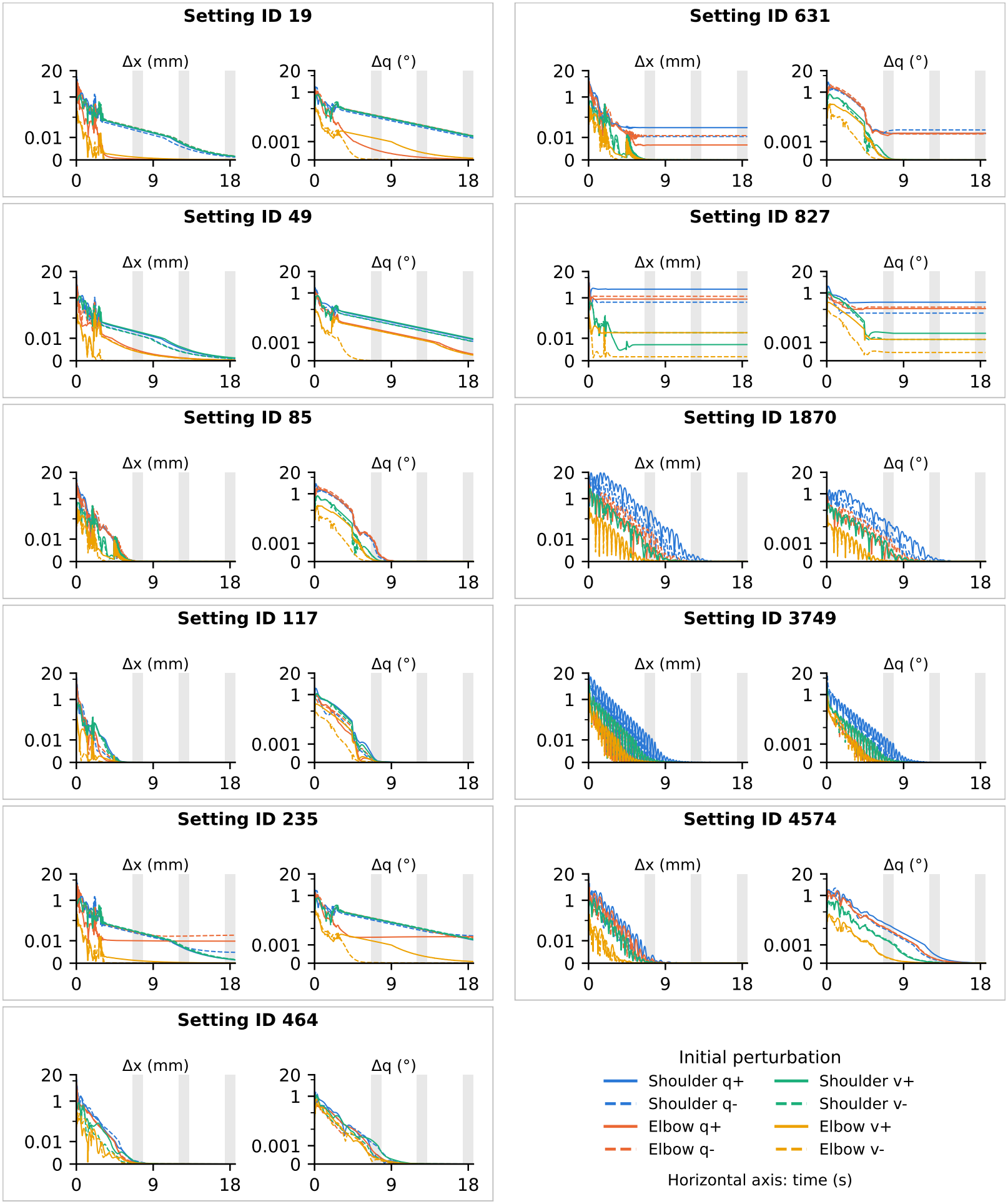
Differences in fingertip position and arm joint angles after an initial perturbation. Each box pairs fingertip distance (left) and joint-angle difference (right) for one setting, plotted against time (s). For each of the 11 controller settings (Table 2), each curve compares a trial starting with a small change in joint angle or velocity to the trial starting from the nominal state. At each saved time, Δ*x* is the distance between the two fingertip positions and Δ*q* is the largest absolute difference between corresponding angles across the seven arm joints. A falling curve means the two trials are becoming more alike in that measure. Colors distinguish changes in initial shoulder or elbow angle (*q*) or velocity (*v*); solid and dashed curves indicate positive and negative changes. Shading marks W1–W3, each containing 60 saved samples. The first sample is one control period after initialization; subsequent samples are spaced by 0.02 s. All settings use the same axes for fingertip distance (0–20 mm) and joint-angle difference (0–20°). The axes are linear near zero (up to 0.01 mm and 0.001°, respectively) and logarithmic above these values.

**Figure S2:**
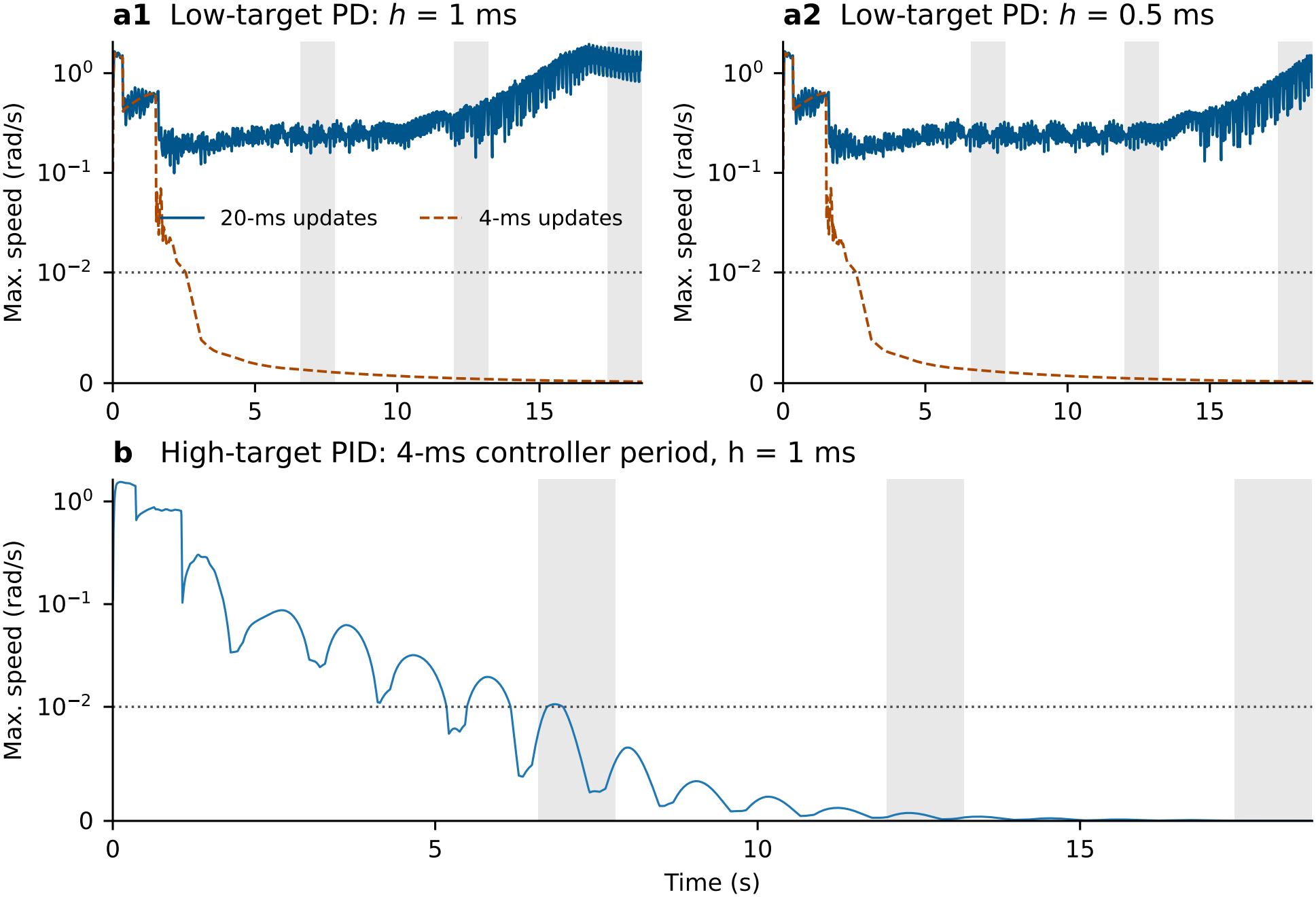
Maximum absolute joint speed in the damped series-elastic variant D08 from the nominal start. **a1, a2** Low-target PD with fixed gains at physics steps of 1 and 0.5 ms, respectively. In both plots, solid lines show 20-ms controller updates and dashed lines show 4-ms updates. **b** High-target PID at its original 4-ms period: speed exceeded the holding limit in the first window only. Shading marks W1–W3; the dotted line is the 0.01-rad/s limit. The speed axis is linear up to that limit and logarithmic above it. Table A1 gives the period-comparison measures.

**Figure S3:**
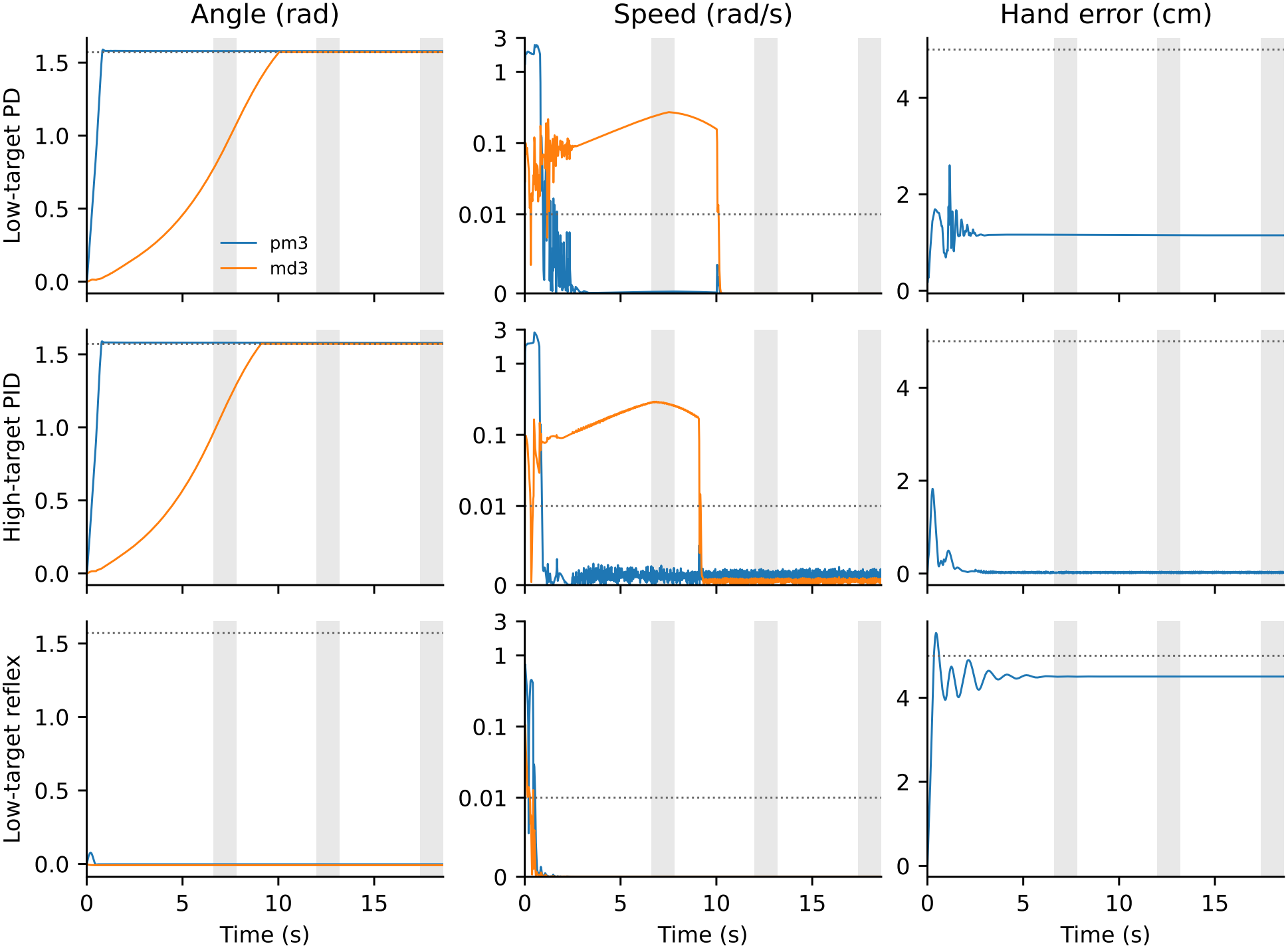
Hand holding and middle-finger motion after transfer to the 63-muscle model. Rows show low-target PD with gravity compensation (setting 85), high-target PID (4574) and low-target reflex (1870) from the nominal start at the 2-ms physics step. Columns show proximal and distal middle-finger flexion (pm3, md3), their absolute speeds and instantaneous index-fingertip error. Dotted lines mark the upper flexion limit (1.571 rad), speed limit (0.01 rad/s) and 5-cm reference for window-mean hand error. Shading marks W1–W3. Traces were saved every 20 ms. Speed is shown on a common scale, linear up to the 0.01 rad/s holding threshold and logarithmic above it. For the high-target PID example, index-finger abduction was fastest in each window. Table A4 reports the full-rate decisions across all joints for all 99 runs.

### A.2 Supplementary methods

#### A.2.1 Endpoint torque and muscle allocation

At each update, the endpoint PD or PID controller converts its requested hand force into a joint torque demand. To choose muscle excitations that can produce that demand, it first calculates how changing each muscle’s activation would change the joint torques at the current joint positions and velocities (*q*, 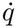).

Let *Q*(*q*, 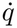, *a*) ∈ ℝ ^20^ denote the vector of joint torques produced by the muscle actuators at joint state (*q*, 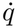) and muscle activation *a* (MuJoCo’s `qfrc_actuator`). At fixed (*q*, 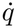), *Q*(*q*, 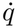, *a*) = *Q*(*q*, 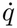, 0) + *S*(*q*, 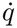) *a*. Each column of *S* gives the change in joint torques when one muscle goes from zero to full activation. At each update the controller computes

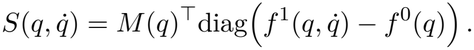

Joint angles *q* determine both the moment-arm matrix *M* (*q*) and the musculotendon lengths. At zero activation, the passive muscle forces *f* ^0^(*q*) depend only on these lengths. At full activation, *f* ^1^(*q*, 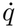) also includes active force, which depends on both musculotendon length and lengthening velocity at the current (*q*, 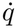).

For each setting, the endpoint controller has a fixed baseline excitation *u*_*b*_. Without gravity compensation, every muscle starts from the same excitation floor *c* when computing its adjustment (*u*_*b,i*_ = *c*). With gravity compensation, the baseline for muscle *i* is *u*_*b,i*_ = max(*u*_0,*i*_, *c*). Initial muscle activation is zero without gravity compensation and *u*_0_ with gravity compensation. At each update it selects an adjustment *δ* and sets the muscle excitation to *u* = *u*_*b*_ + *δ*. To choose this adjustment, it predicts the torque change that would occur if activation reached *u* immediately at the current joint state:

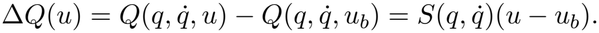

In the simulation, activation follows excitation with 10/40-ms dynamics while the arm continues to move. Across 288 calibration probes, the magnitude of the mean joint torque change over the first control interval ranged from 22% to 88% of the requested magnitude.

#### Independent joint coordinates

Because the 20 joint coordinates are coupled, the endpoint controller expresses both the requested torque and the torque produced by the muscles in the same nine independent coordinates. We denote the angles of the seven independent arm joints and two independent thumb joints by *q*_ind_ ∈ ℝ ^9^ and their angular velocities by 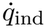. The subscript ind denotes independent coordinates. The matrix *T* (*q*) ∈ ℝ ^20×9^ contains the derivatives of the coupling polynomials at the current *q*; it maps the nine independent joint velocities to all 20 and spans the kinematic tangent of the constraint manifold. Because the constraints are soft, 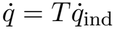 holds only approximately along simulated trajectories; *T* is used only to express demands and forces in the nine independent coordinates. *J* ∈ ℝ ^3×20^ is the translational Jacobian of the fingertip site, *J*_*r*_ = *JT* and *S*_*r*_ = *T*^*⊤*^*S*.

#### Requested hand force and joint torques

The purpose of this calculation is to express the hand-holding task as a torque demand *τ*_req_ at the nine independent joints. This gives the muscle allocation a target: it chooses excitations to approximate the joint torques needed to keep the fingertip near its target and, when posture feedback is enabled, to bias the arm toward the target posture. The PD controller requests a hand force based on the hand’s displacement from the target and its velocity. The PID controller also uses accumulated position error. At update *k*, the controller observes *q*_*k*_,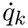 and the fingertip position *x*_*k*_. The target *x*^*\**^ is fixed. The position error is *e*_*k*_ = *x*^*\**^ − *x*_*k*_, and the fingertip velocity is 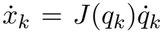. Write *J*_*r,k*_ = *J*_*r*_(*q*_*k*_) and 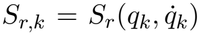. The PID integral *z*_*k*_ accumulates fingertip position error over time (m s). Its update is *z*_*k*_ = *z*_*k* 1_ +*e*_*k*_ Δ*t* unless the anti-windup rule limits or freezes the update. Both PID variants cap the integral contribution in each direction. The conditional variant also retains the previous integral when further accumulation would increase demand in a direction the muscles cannot realize (see Sect. A.2.8 for the update rule). PD uses only the position and velocity terms (*ρ* = 0). With position gain *K*_*p*_ (N/m), integral weight *ρ* (1/s) and velocity weight *τ*_*d*_ (s), the requested hand force and joint torques are

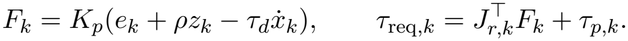

The optional posture torque uses the current independent joint angles and velocities:

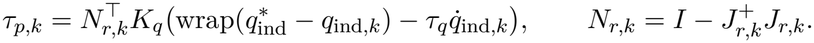

Here 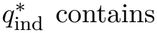 the independent joint angles of the target posture. The function wrap(*θ*) = atan2(sin *θ*, cos *θ*) returns the signed angular error for each joint. The geometric projector *N*_*r*_ projects onto the null space of *J*_*r*_, with *I* the 9 × 9 identity and 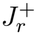 the Moore–Penrose pseudoinverse (relative singular-value cut-off 10^−9^). In settings with posture feedback, the posture gain is *K*_*q*_ = 100 N m/rad and the joint velocity weight is *τ*_*q*_ = 0.1 s. In the other settings, *τ*_*p*_ = 0. The posture term uses the redundancy of the arm to bias the independent joint angles toward their target values. Its projection uses joint directions that leave fingertip position unchanged to first order in the kinematics. The resulting torque need not leave fingertip acceleration unchanged in the simulated dynamics.

#### Muscle excitation

At update *k*, the endpoint controller chooses an adjustment *δ*_*k*_ to the baseline excitation *u*_*b*_, with one component per muscle. For a candidate adjustment *δ*, the model predicts a torque change *S*_*r,k*_*δ*. The controller minimizes the mismatch with the requested torque *τ*_req,*k*_ while keeping each muscle’s excitation between the dimensionless floor *c* and 1:

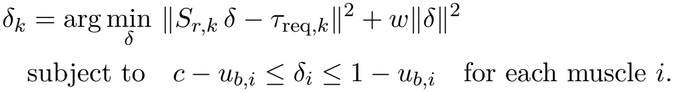

The second term penalizes departures from baseline, with weight *w* = 10^−3^ (N m)^2^. We use bounded-variable least squares (SciPy `lsq_linear` BVLS [19], tolerance 10^−10^, at most 200 iterations). We accept the solution if the solver returns a positive status code and every component of *δ*_*k*_ is finite. If either check fails, the trial is invalid.

The controller adds the accepted adjustment to the baseline and applies the excitation limits elementwise:

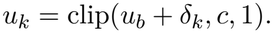

It sends *u*_*k*_ to the muscles in single precision.

#### A.2.2 Paired trajectory measurements

##### Effect of the initial perturbation on the trajectory

For each setting that passed all 27 holding windows, we compared each of its eight perturbed runs with its nominal run at the same elapsed times. This comparison measures how far the trajectories separate after a change in the initial state. At each sampled time *t*, we measured the distance between the two fingertip positions and the largest absolute difference in arm joint angle and angular velocity:

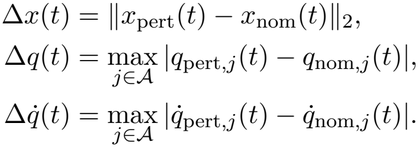

Here *A* contains the seven arm joints (elevation-plane angle, shoulder elevation and rotation, elbow flexion, pronation–supination, wrist deviation and flexion).

The first comparison uses the sample recorded immediately after the first control step (*t* = Δ*t*). Subsequent samples are spaced 0.02 s apart. We calculated differences in double precision from these single-precision samples and summarized their means and maxima within W1–W3. Each window contains 60 samples at either controller period. These differences describe sensitivity to the initial state over the observed interval. Holding was assessed separately using *E, D* and *V* at every control step.

#### A.2.3 Force-map checks and target mechanics

##### Muscle force map and allocation

We tested whether a change in muscle activation produced the joint torque change predicted by *S* and whether the allocated activation produced torque in the requested direction. At the first control step of each endpoint trial, we also compared the analytic *S* with a finite-difference estimate using an activation increment of 0.02 (central differences in the interior, one-sided differences at the bounds). A relative Frobenius difference above 10^−9^ would have stopped the trial.

##### Mechanical sensitivity at the target postures

To characterize how small changes in joint position or velocity affect acceleration at each nominal target, we held muscle activation fixed. With *a* = *u*_0_, the 20 joint angles and their velocities define a 40-dimensional mechanical state. Its 40 × 40 linearization matrix is

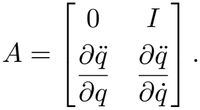

We estimated the two lower blocks by central finite differences of the MuJoCo forward acceleration, which includes the soft equality-constraint forces (widths 10^−4^ rad and 10^−3^ rad/s, repeated with 10^−3^ and 10^−2^). Activation states are not part of *A*. We checked the residual acceleration at each target before interpreting the spectrum. A nonzero residual means that the spectrum describes local mechanical sensitivity at a non-equilibrium state. Equilibrium stability is examined separately in Sect. 2.7.

#### A.2.4 Series elasticity, damping and simulation schedule

##### Series elasticity and damping

We denote the native MuJoCo muscle model by R. Its musculotendon path has no series elasticity. The elastic variants (E02, E04, E08) add a series elastic element to each muscle. The damped variants (D02, D04, D08) add the same elastic element and include series damping. At the reference elastic force *F*_0,*i*_, the series element of muscle *i* has extension *ϵL*_0,*i*_, where *L*_0,*i*_ is the reference length. Thus *ϵ* is a dimensionless extension parameter. The model suffixes 02, 04 and 08 correspond to *ϵ* = 0.02, 0.04 and 0.08, respectively.

The initial screening and long-horizon trials in Sect. 2.4 used a 2-ms physics step. The series element adds an extension state whose force balance is evaluated at each physics step. To resolve changes in extension and tension more finely between controller updates, we halved the physics step to 1 ms. We used the same 1-ms step for the native model R so that the comparison with the six variants used the same integration interval. We retained the controller periods, gains, targets, reference lengths and gravity-compensation excitations. The elastic and damped variants add non-negative series extension *δ*_*i*_ to each muscle while preserving native geometry, inertia, gravity, joint damping, soft constraints and the MuJoCo muscle force law. For each variant, the elastic force, damping coefficient and total series force are

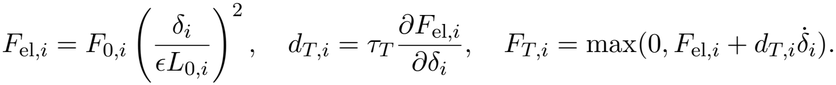

The damping time constant is *τ*_*T*_ = 0 for the elastic variants and *τ*_*T*_ = 0.01 s for the damped variants. To compute muscle force, the MuJoCo muscle model uses the total musculotendon length minus the extension of the series element (*L* _*MT,i*_ − *δ* _*i*_) and the musculotendon velocity minus the extension rate of that element 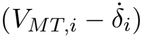. We retained the model’s signed normalization offset. The reference length *L*_0,*i*_ is a model parameter rather than an anatomical tendon length. These are model-sensitivity variants.

##### Integration and initialization

At each physics step, we solved for the extension of the series element in each muscle using a scalar backward-Euler update. We accepted the solution only if it satisfied the model’s criteria for force balance between the muscle and the series element. If the solver could not find an acceptable solution, we stopped the simulation and recorded it separately from trials that reached the end but failed the holding criterion. We did not clip the extension rate to continue a stopped trial. Muscle activation followed the native explicit update. Endpoint controllers retain their rigid static surrogate, and the reflex senses musculotendon length and velocity. These are transfer tests without retuning.

For initialization, reflex and settings without gravity compensation start with *a* = 0, while settings with gravity compensation start with the nominal *u*_0_. A positive tonic floor does not replace this initial activation. Position/velocity offsets retain the original activation and controller memory; we initialized each series element at the smallest extension satisfying static force balance at the prescribed joint position and original activation, including for velocity-offset starts. Dynamic equilibrium at initialization is not asserted.

##### Settings and initial conditions

The main sensitivity schedule has 429 cases: R/E04/D04/D08 each use 11 settings and nine starts; E02/E08/D02 use only the 11 nominal starts. To check sensitivity to the physics time step, we halved it to *h* = 0.5 ms for six predefined runs: the low-target PD with gravity compensation, middle-target reflex and high-target PID settings in each of the E04 and D04 models. These six runs and the additional diagnostic runs were separate from the 429 main cases. Holding is assessed across all three windows.

#### A.2.5 Feedback timing and excitation interventions

After examining the holding trials, we selected D08 for post-hoc tests. These runs used the nominal start or states reached from that start.

##### Controller update period

For the PD controller, we compared controller periods of 20 and 4 ms at unchanged gains in R, D04 and D08, using a 1-ms physics step. D08 at 4 ms was also tested at a 0.5-ms physics step, with the existing 20-ms diagnostic at that same step as its comparator. Each run lasted 18.6 s. We assessed holding at every control update. To quantify joint-velocity fluctuations, we sampled each joint’s velocity every 20 ms over (6.6, 18.6] s and computed its root-mean-square (RMS) deviation from its own interval mean. We compared two measures: the RMS fluctuation in wrist deviation velocity and the largest RMS fluctuation among the seven arm joint velocities. If either measure in the 20-ms baseline trial was at most 10^−4^ rad/s, we omitted both ratios from the comparison. Otherwise, for each measure, we divided the value from the 4-ms controller trial by the value from the 20-ms controller trial with the same physics time step. We classified the reduction as large if both ratios were 0.20 or less. The two-step comparison required the 4-ms amplitudes at the two physics steps to agree within the larger of 20% and 10^−4^ rad/s. These are operational finite-time criteria, not stability tests.

##### Timing of muscle excitation at a fixed update period

To test how the timing of muscle excitation affected ongoing oscillations in D08, we compared three simulations starting from the same state at 6.6 s. In the reference simulation, the PD controller continued to update every 20 ms. In the other two, it computed a new excitation every 4 ms. We applied each new excitation to the model muscles immediately in one simulation and 8 ms after computation in the other.

We chose 8 ms to match the average time since the applied excitation was computed. When an excitation is applied immediately and held until the next update, this time increases from zero to one update period and averages half that period (10 ms for 20-ms updates and 2 ms for 4-ms updates). Adding 8 ms to the latter gives the same 10-ms average with more frequent updates. This timing comparison was independent of physiological nerve conduction times.

Each simulation continued for 13 s with unchanged gains. As in the preceding test, we ran the comparison at physics time steps of 1 and 0.5 ms, using the state saved at the corresponding step size. We compared the slopes of log oscillation amplitude over (0.15, 0.60] s after branching to assess whether the delayed 4-ms response was closer to the immediate 4-ms response or to the 20-ms reference (Sect. A.2.9).

##### Halving or removing fluctuations in muscle excitation

We tested whether arm oscillations changed when we reduced the variation in each muscle’s excitation around its recent mean. We started from states saved at 6.2 and 6.7 s in the D08 baseline run and at 6.2 s in D04. At each starting point, we averaged the preceding 50 excitation commands for each muscle (1 s at the 20-ms update period) and kept this reference vector *ū* fixed. At each subsequent update, *u*_fb_ denotes the vector of muscle excitations that the PD controller would ordinarily apply at the current state. We used this vector to compute the excitation *u* actually applied to the muscles,

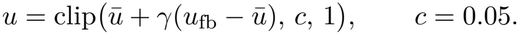

In D08, we retained the controller output (*γ* = 1), halved each muscle’s deviation from its reference (*γ* = 0.5), or held each excitation at its reference (*γ* = 0). In D04, we compared only the unchanged and constant excitations. Muscle activation evolved normally.

For each starting point, we first verified that the unchanged controller reproduced the saved baseline trajectory over the next 4 s. We planned to compare motion amplitudes in the final second, (3, 4] s after the start, provided that the trajectories remained close. At each physics step, we compared each intervention with its baseline at the same elapsed time. We stopped the intervention if any of the 20 joint coordinates differed by more than 0.05 rad or if the fingertip positions differed by more than 0.01 m. An early exit therefore prevented the planned amplitude comparison.

##### Reducing excitation fluctuations by 5% or 10%

In a post-hoc test, we asked whether smaller reductions in excitation fluctuations changed the amplitude or frequency of arm oscillations in D08. We selected four starting points at different phases of a single oscillating trajectory (PD controller period 20 ms, physics time step 1 ms). From each starting point, we ran four conditions for 1.8 s each. For the first 0.8 s, one condition used the ordinary controller output. Two others reduced each muscle’s deviation from its pre-branch mean by 10% (*γ* = 0.90) or 5% (*γ* = 0.95).

Reducing these deviations could also shift the mean excitation during the measurement window. To test the effect of this shift alone, the fourth condition added a constant offset to each muscle’s ordinary excitation. We chose the offsets before collecting intervention data to match the mean change predicted for each muscle under the 10% reduction.

We measured motion from 0.4 to 0.8 s after each start by fitting a sinusoid and a linear trend to projected arm velocity. One fit used the pre-branch frequency (7.33 Hz) to compare amplitudes at a common frequency. A second fit estimated frequency separately for each trace. For the 10% reduction and its constant-offset control, we checked how closely the realized mean excitation changes matched and whether the offset control introduced excitation fluctuations in the oscillation band (D1). We also checked whether cycle-averaged shifts in joint angles and fingertip position agreed between the two conditions to within half a baseline oscillation amplitude (D2). These 16 continuations were separate from the 16 runs in the initial protocol (update-period comparison, strong excitation reductions and integral-action test). Six of those initial runs checked reproducibility: three baseline replays and three restarts with *γ* = 1.

##### Did integral action change muscle excitation?

In D08, we compared the PID controller’s joint-velocity fluctuations at 4-ms updates with its own 20-ms baseline, using a 1-ms physics time step. To determine whether integral action contributed at 4 ms, we tracked the integral state and recorded when the anti-windup rule prevented it from updating. We then compared the excitation applied to each muscle and the resulting trajectory with those of the PD controller at the same 4-ms update period.

#### A.2.6 Closed-loop fixed points and branch classification

The state includes all 20 joint positions and velocities, 34 muscle activations and 20 solver-warmstart entries. For PID, the map also includes the integral of fingertip position error, *z*_*k*_. Series variants add 34 extension states. Native and qualified float64-command reference maps are separate. Full-state defects and speed, acceleration and activation-derivative tolerances qualify candidate fixed points. Activation/deactivation-side extensions and a bounded number of mixed extensions are evaluated with base-map, branch/domain and three-width convergence gates. We recorded allocation masks, clipping, constraints and native command quantization. For PID, we also recorded how the anti-windup rule treated the integral update, including whether the conditional rule retained the previous integral. A failed finite-width gate does not prove that a classical derivative is absent in every smaller neighbourhood.

Protocol A0 examined the rigid model, and protocol A1 examined the series-elastic models. A1-R denotes a post-hoc reanalysis of the branch patterns identified in A1, using the same models and the same reference map. We reclassified those patterns using the number of force-equilibrium roots, the selected root and the retained solver features. We recorded solver origin and bracket indices separately for reference (see Sect. A.2.12 for the A1 and A1-R comparison).

#### A.2.7 Full-state perturbations, response matrices and validation

We used four solver settings (L0–L3). From L0 to L3 the iteration limit doubled at each level (100 to 800), and the tolerance decreased tenfold (10^−8^ to 10^−11^). For a reference state *Y*_*\**_ tested against Φ(*Y* ^*\**^) = *Y*_*\**_, we prescribed *Y*_0_ = *Y*_*\**_ + *rDd*, where *d* specifies the perturbation direction and *r* ∈ {4, 8, 16} its amplitude multiplier. The diagonal entries of *D* specify the base sizes: 10^−4^ rad for joint position, 10^−3^ rad/s for velocity, 10^−3^ for activation, 10^−2^ rad/s^2^ for solver warmstart and 10^−4^ m s for the PID integral state. At update *k*, the measured response was

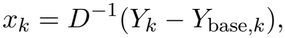

relative to an unperturbed run with the same feedback controller and solver setting. The physical/controller block contains positions, velocities, activations and, for PID, the integral state. The warmstart block contains the remaining numerical coordinates.

At each solver setting and amplitude we estimated a one-update map 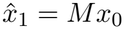. To estimate column *j*, we perturbed only initial coordinate *j* by +*rD*_*jj*_ or − *rD*_*jj*_ as permitted. The corresponding full response vectors after one update are 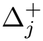 and 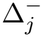. When both signs were feasible for every coordinate, we used

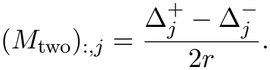

If an activation bound ruled out one perturbation sign for any coordinate, we constructed the columns of *M*_bound_ as follows:

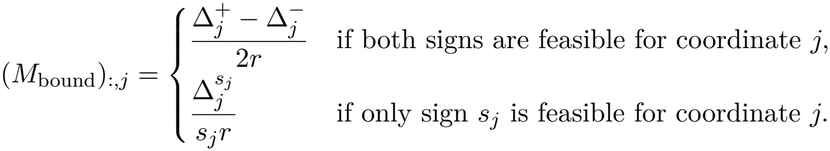

Here *s*_*j*_ = +1 for a permitted increase and *s*_*j*_ = −1 for a permitted decrease. Predictions with *M*_bound_ were restricted to directions allowed by the activation bounds.

A missing required response invalidated the map.

We fixed three prediction tests before acquisition. First, matrices at adjacent amplitudes (16 versus 8 and 8 versus 4) had to agree within 5% in both relative operator and Frobenius norms for the full matrix and both diagonal blocks. Second, we tested one-update predictions in 16 independent mixed-coordinate directions per solver level and amplitude. Third, we applied the same matrix repeatedly and evaluated the predictions against the matched baseline at 0.2 and 2 s, flagging exits from the feasible cone. For either block *B*, a resolved response had to satisfy

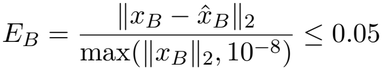

and the prescribed component-wise bound. Small responses were assessed against absolute precision floors derived from baseline reproducibility and eight fixed perturbation directions at unit-in-the-last-place precision; short and long trajectories had to agree over their common prefix. The 16 numerical and completeness checks were separate from predictive validity. Controls perturbing warmstart or disabling it ran for 0.2 s. Section A.2.10 reports the results.

#### A.2.8 PID integral update

Before the first PID update (*k* = 0), the stored integral is initialized to *z*_−1_ = 0. At update *k, e*_*k*_ denotes the current fingertip position error and *z*_*k*−1_ the integral state before the update. The controller computes a trial integral and command, then applies the anti-windup rule chosen for that setting:

- Trial integral: 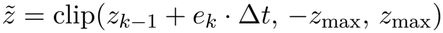 componentwise, with *z*_max_ = 40 N*/*(*K*_*p*_*ρ*), so that 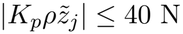.
- Trial command: evaluate the force, torque and allocation equations at the same arm state using 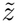 as the integral value, yielding 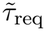and *ũ*.
- *Clamp* anti-windup: 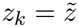 and *u*_*k*_ = *ũ*.
- *Conditional* anti-windup: with the change in integral generalized force 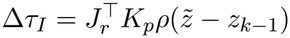 and the unrealized trial demand 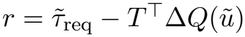, the integrator is frozen if *∥* Δ*τ*_*I*_*∥* > 10^−12^, *∥ r∥* > 10^−12^ and cos(Δ*τ*_*I*_, *r*) > 10^−6^. Then *z*_*k*_ = *z*_*k*_ −_1_ and the allocation is solved again, using the retained integral value to obtain *u*_*k*_; otherwise 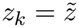 and *u*_*k*_ = *ũ*.

#### A.2.9 Command-delay comparison

For the first 8 ms after the switch, the delayed controller continued to apply the last excitation from the 20-ms run. We verified that the immediate 4-ms implementation reproduced the saved 4-ms continuation over the first 25 updates. The two delayed runs were counted separately from the 16-row modulation protocol and excluded from the holding counts. We projected the physics-rate joint positions onto the first direction of the 5–10 Hz band in the saved late-window trace for PD control in D08. A least-squares fit of a constant, a linear trend and sine and cosine terms at 7 Hz over one-cycle windows (quarter-window hop) gave a local amplitude estimate. We regressed log amplitude on window-centre time in (0.15, 0.60] s after the switch. For each physics step and estimator, the contrast was *R* = (*σ*_delay_ − *σ*_4_)*/*(*σ*_20_ − *σ*_4_). The reference-rate difference had to be at least 4 s^−1^, and all amplitudes in the window had to be at least 100 times the mean of the corresponding 4-ms sham over relative (1, 12] s. The 20-ms-side classification required 0.75 *≤ R ≤* 1.25; agreement across step sizes and consistency with two secondary envelopes (causal bandpass RMS and the seven-arm-joint speed envelope) were checked. In the delayed runs, the latter envelope averaged 7.17 and 7.21 rad/s over relative (9, 12] s, compared with 1.65 and 0.92 rad/s in the 20-ms continuations (rounded), in order of decreasing physics step. These late amplitudes were descriptive and did not enter the early classification.

#### A.2.10 Full-state response results

Under the criteria in Sect. A.2.7, every setting failed at least one adjacent-amplitude comparison (Table A2).

The primary qualified signed rows number 228 for the low-target PD setting and 384 for each other setting; rows for the two maps are not pooled or counted twice. For that setting, 156 additional rows fell outside the feasible perturbation cone and were excluded from the prediction test. No qualified row passed the joint prediction criterion. The physical-error medians at multipliers 16/8/4 were 0.556/0.553/0.574 (low-target PD), 0.211/0.135/0.135 (middle-target reflex), 0.339/0.360/0.346 (high-target PID), and 0.123/0.117/0.117 (low-target reflex). The L0 physical-column classifications include respectively 35, 1, 14 and 3 scale-dependent columns; global positive homogeneity is therefore not claimed. Matrix radii, qualified-row errors and the distinction between the two maps are reported in Table A2.

#### A.2.11 Prospective singular-direction protocol

We used the L0 map at *r* = 8 and the state scaling defined in Sect. A.2.7. Let *M*_phys_ denote its submatrix with physical/controller coordinates as both inputs and outputs. For an observation time *k* updates after perturbation, we formed (*M*_phys_)^*k*^ and retained the input columns for joint positions, joint velocities and muscle activations. The leading right singular vector of this matrix specifies how to perturb these coordinates together to maximize the predicted response norm for a unit input in scaled coordinates [23]. In the simulations we applied this direction with the sign allowed by the activation bounds for PD and with both signs for PID. Initial solver warmstart values and the PID integral state were unchanged.

For each tested sign, we measured the gain

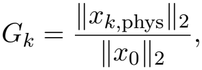

where the numerator is the physical/controller state difference from the unperturbed run at the selected time and the denominator is the measured initial perturbation size. At *r* = 8, we compared this gain with two benchmarks at the same time and in the same setting. The first was the largest gain from four earlier runs using two fixed perturbation patterns, each tested with both signs. The second was the largest gain from new runs using one fixed pseudorandom direction (one sign for PD, both for PID). A response passed if its gain was at least 1.25 times each benchmark. At *r* = 4, we compared gains only with the earlier runs at that amplitude. Before the new runs, we fixed the directions and their orientation, thresholds, eligibility rules and acquisition schedule. The two saved coordinate patterns were all joint positions positive and alternating joint velocities. The pseudorandom direction was a normalized standard-normal 74-vector with a seed fixed before acquisition; its smallest acute angle to every candidate and saved direction was 78.8°. For the low-target PD setting, candidate signs were restricted to the one-sided activation cone, and predictions were labelled as extrapolated from the first transition whose input left the cone (update 4 for the one-update and 0.2-s directions, update 69 for the 2-s direction). A new run was eligible at update *k* only under the following conditions:

- its baseline reproduced the saved baseline;
- the restored initial state was read back bitwise;
- the L0 precision checks passed up to *k*;
- it had not left the domain at or before *k*;
- its initial warmstart and integral-state differences were zero;
- *∥x*_*k*,phys_*∥*_2_ was at least 100 times the precision floor at update *k*.

Runs stopped by a non-finite state were to be ineligible at every time; none occurred. A four-run reproducibility check preceded the 23 prospective runs and reproduced the saved baselines and initial states. The four-level point check of the saved protocol was not applied to new runs.

#### A.2.12 Series-model and local-diagnostic qualifications

##### Static stiffness and mechanical perturbations

Let *F* be the external force applied to the hand, *x* its position in world coordinates and *g*(*q, F*) the joint acceleration at zero velocity and fixed activation. At equilibrium, *g*(*q, F*) = 0. Differentiating this condition gives the hand compliance matrix

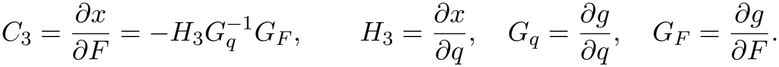

For a horizontal force increment, the upper-left 2 × 2 block *C*_*xy*_ maps the applied force increment to the horizontal displacement. Its inverse 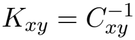 is the static horizontal hand stiffness.

We checked this estimate by independently solving for the equilibrium displacements under small horizontal loads. We tested agreement for additional force directions and at smaller loads, checked that residual errors were small relative to the measured displacements, and repeated the measurement with added springs of known isotropic and anisotropic stiffness. Positive definiteness was not required.

For the separate post-hoc constant-excitation test (C0-S) at the middle and high targets, we linearized the dynamics of the 20 joint positions *q* and 20 joint velocities *v* about each equilibrium. Using central differences, we estimated the Jacobian of (*v*, 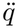) with respect to (*q, v*). We selected the eigenvalue with the largest real part. Its eigenvector specified a direction in which to perturb both positions and velocities; a positive real part predicts local growth along that mode.

To test this prediction in the simulator, we scaled the eigenvector so that the largest initial change in any joint angle was 10^−6^ rad, scaling its velocity components by the same factor. At each of two physics time steps (1 and 0.5 ms), we ran three trajectories for 0.5 s: one from the equilibrium and one from each of the two opposite perturbations. We measured how the joint-position difference from the unperturbed trajectory changed over that interval and verified at each step that muscle activation remained constant. The convergence and constraint checks are described below.

R/E04/D04/D08 have nine starts per setting, while E02/E08/D02 have only nominal starts. The six predefined half-step checks are five completed holds and one E04 domain stop, separate from the main 429 cases and additional diagnostics. The strict serial equilibrium rule and native-normalized extension are disclosed sensitivity assumptions. No parameter or initial-state rescue is used after a root failure.

A0 has 14 rigid protocol rows: 11 N2 rows at 2 ms and three N1 rows at 1 ms. Of these, 11 are allocation rows and three reflex rows; the seven reference-root-verified allocation rows are a subset. A1 has six planned elastic rows; E04/3749 inherits its source stop. Native fixed-point verification in 10 rigid rows includes h-specific repeats. A finite stencil gate failure does not prove mathematical non-differentiability in every smaller neighbourhood. Extension spectra do not establish the stability of the switching native map.

The elastic A1 reference classification yielded 18 patterns (17 branch boundaries and one extension/base mismatch). A1-R classified the same patterns without using the bracketing origin as a criterion. Of the 18 patterns, eight still crossed a branch boundary, eight left the specified state bounds, one extension missed the verified fixed point and one lacked a continuous series-equilibrium solution. Some B1 domain-stop subtypes were also descriptive post-hoc annotations, without changing primary status or the holding population.

The controller-state defect tolerances are 10^−8^ rad for position, 10^−7^ rad/s for velocity, 10^−8^ for activation, 10^−6^ rad/s^2^ for solver warmstart, 10^−9^ m s for integral state and 10^−8^*L*_0_ for extension. Additional base gates are maximum speed 10^−6^ rad/s, acceleration 10^−4^ rad/s^2^, activation derivative 10^−6^ s^−1^, extension rate 10^−6^*L*_0_/s and hand error below 0.05 m. The controller-map FD base widths are position 10^−4^ rad, velocity 10^−3^ rad/s, activation 10^−3^, warmstart 10^−2^ rad/s^2^, integral state 10^−4^ m s and extension 10^−4^*L*_0_, each at multipliers 1, 0.1, 0.01. Scaled-map adjacent-width differences must be at most 0.01 and spectral-radius differences at most 0.001. No failed width is silently replaced with a narrower one. Native float32 commands and the qualified float64 reference are separate maps.

The C0 stiffness pilot gates required maximum acceleration 10^−4^ rad/s^2^, series-force residual 10^−6^*F*_0_, seed position change at most 0.05 rad and hand displacement at most 1 mm. C-R was designed after the C0 pilot. It uses saved rigid static bases and the original holding activation, without optimizing activation or changing the plant/solver to obtain a pass. Central position widths are 10^−5^, 10^−6^, 10^−7^ rad, paired with force widths 10^−4^, 10^−5^, 10^−6^ N. Three-width compliance/stiffness differences must be at most 1%, with scaled *G*_*q*_ condition number at most 10^10^, compliance condition number at most 10^4^ and solve residual at most 10^−8^. The smallest declared width is the representative measurement.

Known springs used *κ* = max(1, 2 max |*λ*(sym*K*_*xy*_)) | N/m; *κ* > 10^4^ stopped the measurement instead of clipping it. The isotropic spring was *κI* and the anisotropic spring was a 30-degree rotation of diag(2*κ, κ*), both anchored to the original base. A common probe force was selected before the probes from 10^−5^, 3 × 10^−5^, 10^−4^, 3 × 10^−4^, 10^−3^, 3 × 10^−3^, 10^−2^ N, using all three contexts: no added spring, isotropic and anisotropic known spring. The selected maxima were 3 × 10^−4^, 3 × 10^−3^ and 10^−3^ N for the low-target PD, middle-target reflex and high-target PID settings. Selection required predicted 3D displacement at most 0.25 mm, preserved joint-limit margins and predicted xy signal at least 10^−8^ m. A zero-force verification root in each context supplied the displacement origin. Its drift and the local residual indicator 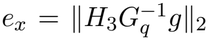 had to be at most 1% of the minimum signal, using predicted signals at force selection and actual signals after probing. This indicator is not a strict error bound; force-induced acceleration ratios were recorded for reference in C-R. Each context used 24 training and eight independent-direction holdout probes, with every root started from its base and without using the IFT derivative as the root solver’s Jacobian or initial displacement. Actual roots had to preserve the active branch, stay within the 1-mm/0.05-rad domain, resolve xy displacement above 10^−8^ m and satisfy the C0 acceleration tolerance together with *e*_*x*_ *≤* 0.01 *∥*Δ*x*_*xy*_*∥* _2_. The same-context representative *G*_*q*_, *H*_3_ were used for this diagnostic, not fed to the independent root solver.

Zero-intercept force/compliance fits required IFT/probe agreement at most 5%, *R*^2^ *≥* 0.98 with the sum of squares about the sample mean, holdout relative error at most 5% and 5% low-amplitude consistency. Each method had to recover both known isotropic and anisotropic spring additions within 5%. The logical conjunction of base, branch, width, conditioning, resolution, fit, holdout and spring checks defines instrument validity; positive definiteness is not a criterion. Three settings passed (Table A3). Only one nominal static base per target was tested; no multiple-start or elastic stiffness validation is inferred.

For C0-S, central differences used position and velocity increments of 10^−4^ rad and 10^−3^ rad/s, scaled by 1, 0.1 and 0.01. We scaled the Jacobian by the base increments and checked constraint branches. Adjacent relative Frobenius differences had to be at most 0.01; leading real parts had to agree within the larger of 10^−3^ s^−1^ and 1% of the larger absolute leading real part. We checked activation byte for byte without overwriting it. State and fingertip records used post-step times and constraint and solver records used pre-step times. Comparisons after a branch change were excluded by design.

C0-S was specified after the stiffness pilot. The reported analysis used 9000 physics steps.

#### A.2.13 Detailed perturbation and stiffness results

##### Effect of added command delay

For the PD controller in D08, the early log-amplitude slopes for the 4-ms switch, the same switch with 8 ms added delay, and the 20-ms continuation were − 6.790, +0.498, and +0.309 s^−1^ at *h* = 1 ms, and − 6.881, +0.730, and − 0.292 s^−1^ at *h* = 0.5 ms. The normalized contrasts were 1.027 and 1.155, both in the prespecified 20-ms-side interval of 0.75–1.25. Both secondary measures gave the same classification at both physics steps. The late motion in the two delayed runs was larger than in the corresponding 20-ms continuations (Sect. A.2.9).

##### Mean-command probe

All three *γ* = 1 arms reproduced their 4-s saved baseline tails byte for byte. All five *γ* < 1 arms exited the prespecified nearby comparison domain before reaching the primary interval (3–4 s after the intervention began). In D08, *γ* = 0 exited through the hand criterion at 0.195 and 0.104 s; *γ* = 0.5 exited through the joint criterion at 0.612 and 0.556 s. D04 *γ* = 0 exited through the joint criterion at 2.750 s, at 0.050036 rad. Because neither strong intervention level reached the primary interval within the nearby domain, the effect of reducing state-dependent command modulation on the sustained motion remained inconclusive. These were comparison-domain exits, not tests establishing divergence, instability or failure to suppress velocity fluctuations.

##### Weak command modulation

All 16 continuations completed within the comparison domain. At 0.90, the four fixed-frequency amplitude ratios ranged from 0.580 to 0.745 and the free-frequency ratios from 0.593 to 0.747. Fitted frequencies fell by 3.9–6.7%. Constant-offset controls yielded amplitude ratios of 0.991–1.003 and passed the live command checks at all four phases. Fingertip shifts met the cycle-averaged shift criterion (D2) at all four phases. Joint shifts exceeded the same tolerance at every phase, with maximum differences of 9.6–23.1 mrad. The amplitude reduction met the prespecified criterion at all four phases, while the mechanism assessment was reserved at each phase. For the exploratory 0.95 condition, fixed-frequency ratios were 0.809, 0.831, 0.658 and 1.104.

##### Conditional PID and duplicate trajectory

The conditional PID controller at 4 ms held in all three windows in D08. Its deviation and maximum-arm RMS fluctuation ratios relative to its own 20-ms baseline were 0.001357 and 0.001094. However, the accepted integral state remained zero: the initial error was zero, and conditional anti-windup froze the update on each of the remaining 4649 control instants. The final commands and state trajectory were byte-identical to those of the PD controller at 4 ms across all 54 shared saved fields. This is consistent with the conditional anti-windup freezing reported in the existing controller comparison (Sect. 3.5) and with the identical nominal command sequences of settings 631 and 85 in the native runs (Sect. 3.4). This configured-PID result therefore provides no distinct evidence for a contribution of integral action. It is a property of this run, not general controller equivalence. In the 20-ms baseline, the integral state changed on five control instants and the command stream first differed from setting 85 at index 28.

##### Controller-state qualification

Native fixed-point defects passed in 10 rigid protocol rows (including repeated settings at different physics steps) and four elastic rows. In the rigid and series-elastic fixed-point analyses (A0 and A1) and the post-hoc reclassification of the elastic results (A1-R), no qualifying native full-state spectrum was obtained. Across the 20 protocol rows, the checks distinguished failure to verify a fixed point from failures of precision, base-map agreement, branch consistency or domain validity. Those protocols exported only specified reference extensions of rigid reflex setting 1870; their expanding modes were not consistently realized by native probes and are not evidence of native instability.

##### Native full-state response and prediction

We tested whether a linear map estimated from perturbations of one amplitude could predict the response at another amplitude or in a new direction. The subsequent four-setting protocol completed all 204416 new updates and 899120 physics substeps, including its acquisition prefixes, with 7400 saved N0 updates reused separately. All 16 numerical and completeness gates passed in every setting: all required records and matrices were available, physical and warmstart precision checks passed, baseline and warm-off responses reproduced, all 9904 applicable initial-update comparisons agreed, and all 32 short/long shadow pairs agreed. No new trajectory left its declared domain. The 9904 comparisons include 2504 on new native records; the rest concern reused N0 records and are not independent new simulations.

Every setting failed at least one adjacent-amplitude operator-consistency test (Table A2). At all four solver levels, the low-target PD setting passed 16/8 but failed 8/4; both reflex settings failed 16/8 but passed 8/4; the high-target PID setting failed both pairs. Thus the operators themselves depend on the tested amplitude, even where mixed-input error medians vary little. None of the qualified mixed-input rows passed the joint linear prediction test either. The physical-state relative errors ranged from 0.059 to 1.461, and the repeated-update predictions also failed the joint criterion. The primary operators were *M*_bound_ for low-target PD and middle-target reflex, and *M*_two_ for high-target PID and low-target reflex. Their L0/L3 spectral radii were 0.9715–0.9989, but these are finite-amplitude secant/cone operators whose predictive validity failed. They are not validated Jacobian stability spectra. This is a complete negative prediction result rather than missing numerical evidence, and does not negate the finite-time holding examples.

##### Prospective singular-direction responses

The prospective protocol completed all 23 scheduled runs (8300 updates, 46000 physics substeps). Both baselines reproduced the saved baselines in every recorded field, and no run left its domain. At *r* = 8, the candidate gain exceeded the maximum of both comparison groups in all nine comparisons, and 7/9 met the predeclared 25% margin. All three one-update comparisons passed. At 0.2 s and 2 s, 4/6 passed. The low-target PD setting passed at both times (gains 19.26 and 2.99; ratios to the saved group 1.49 and 2.48). The high-target PID setting passed for one sign at each time and fell short for the other: the negative sign at 0.2 s (ratio 1.09 to the saved group) and the positive sign at 2 s (1.01). Ratios to the pseudorandom group were at least 3.92 in all nine comparisons. The gains quantify amplification in the *D*-scaled state norm. Cross-setting comparisons use the common physical times of 0.2 s and 2 s. At all 63 L0 evaluation points (21 perturbed runs × three times), neither the physical/controller nor the warmstart block met the full-state response error criterion. The auxiliary reduced physical/controller criterion (relative gain error *≤* 0.05 and cosine *≥* 0.95) passed at 0/6 selected-time points for PD and at 7/12 for PID. Twelve evaluation points for PD, including both its 0.2-s and 2-s selected times, are extrapolations beyond the prediction cone. The four-level point check was unavailable for these L0-only runs.

At the states tested in the muscle-force diagnostic, the model actuator LAT1 had zero active and passive muscle forces because its normalized length is outside the force–length support. Its compiled lengthrange matches an explicit value in the installed XML. This explains the zero activation-sensitivity column at those states and is consistent with zero series extension. It is a model disclosure, not a statement about every configuration or human anatomy.

##### Constant-excitation mechanics

The initial C0 stiffness-pilot bases passed the residual criterion, but none passed all instrument gates: the low-target PD setting crossed a wrist-limit active set in its position stencil, the middle-target reflex setting exceeded the 1-mm displacement domain in an isotropic-spring probe, and the high-target PID setting had no supported force scale.

The C-R protocol passed all gates in three rigid settings, one at each target (Table A3). The eigenvalues of sym(*K*_*xy*_) were 2.751/59.690 N/m at the low target, − 19.573/39.925 N/m at the middle target, and − 6.248/6.735 N/m at the high target. Thus the stiffness matrix was positive definite at the low target and indefinite at the other two; indefiniteness was a measurement result, not a failed validation. Relative errors between the implicit-function-theorem (IFT) estimate and the force probes were 4.85 × 10^−6^, 6.33 × 10^−6^ and 1.11 × 10^−5^. Base-context holdout relative errors were 0.00175, 0.00110 and 0.00114; all known-spring relative errors were below 8 × 10^−6^.

These are fixed-activation, same-active-branch static operators in world *xy* with *z* free. They include muscle, gravity/bias and soft-constraint effects and are not muscle-only or feedback-controlled stiffness. Along the unit minimum-eigenvalue direction of the total matrix, actuator/bias projections were − 18.499/ − 1.075 N/m at the middle target and − 1.254/ − 5.046 N/m at the high target. These projections use the same total direction, not sums of component eigenvalues or a unique causal decomposition. At the low target, the larger-stiffness eigenvector is at 66.0 degrees from world +*x* toward +*y* (modulo 180 degrees), and the larger/smaller stiffness-eigenvalue ratio is 21.7. These are model-coordinate quantities, not anatomical orientation or human-stiffness comparisons. Human postural stiffness has been measured from restoring forces during imposed hand displacements [21]; our calculation keeps muscle activation fixed. No ellipse is drawn for the indefinite cases.

With excitation held at the holding activation and feedback removed, the 40-dimensional mechanical linearizations at the middle and high targets had leading real eigenvalues 4.084 and 1.219 s^−1^ (e-folding times about 0.25 and 0.82 s). Both passed three-width convergence. The leading directions mainly involve forearm rotation and elbow motion. In 0.5 s, joint-position differences from baseline increased by *factors of* approximately 7.71 and 1.84, with both perturbation signs and both physics steps (Fig. 6). Activation remained byte-identical and no constraint active-set change was observed. In these two settings, velocity effects substantially lower the rates relative to position-only estimates. Setting the acceleration derivative with respect to velocity to zero in the same mechanical matrices gave leading rates of 20.29 and 6.99 s^−1^ at the middle and high targets, respectively. At the middle and high targets, the C-R measurement and C0-S test reuse the same saved static joint-position base and holding-activation capture. Their indefinite static xy stiffness and expanding mechanical modes are complementary observations at those bases, not equivalent tests: the former leaves *z* free in a static projection, whereas the latter retains the 40-dimensional position/velocity dynamics. These are numerical local open-loop instability results on the invariant *a* = *a*_*h*_ set; activation perturbations and elastic settings were not tested. Feedback was disabled in these tests.

##### Search for distinct equilibria at the same target

The static stiffness measurement passed in three rigid, fixed-activation states, with one base per target. In saved trajectories for those settings, the nine tested starts per setting converged to the same endpoint within the recorded precision: at 18.6 s the maximum pairwise differences were 1.7 × 10^−7^ rad in joint-position norm and 6 × 10^−8^ in activation norm. All endpoints were within 1.1 × 10^−6^ rad in joint-position norm of their corresponding stiffness base. This comparison uses saved states. In a separate 1-ms test of setting 3749 at R_mid, eight starts used displacements with nominal norms of 0.10 or 0.20 rad in five arm coordinates, in both senses along two hand-position-preserving directions. All eight starts passed all three holding windows. All eight fixed-activation roots qualified and matched the existing stiffness base within the prescribed joint-angle and activation tolerances (10^−5^ rad and 10^−6^ in maximum absolute difference, respectively). Their hand positions were within 1 mm of the existing base. The maximum absolute joint-angle difference from the existing base was 1.1 × 10^−7^ rad. The distinct-base count was zero, so no additional stiffness was measured.

### A.3 Supplementary tables

**Table A1:**
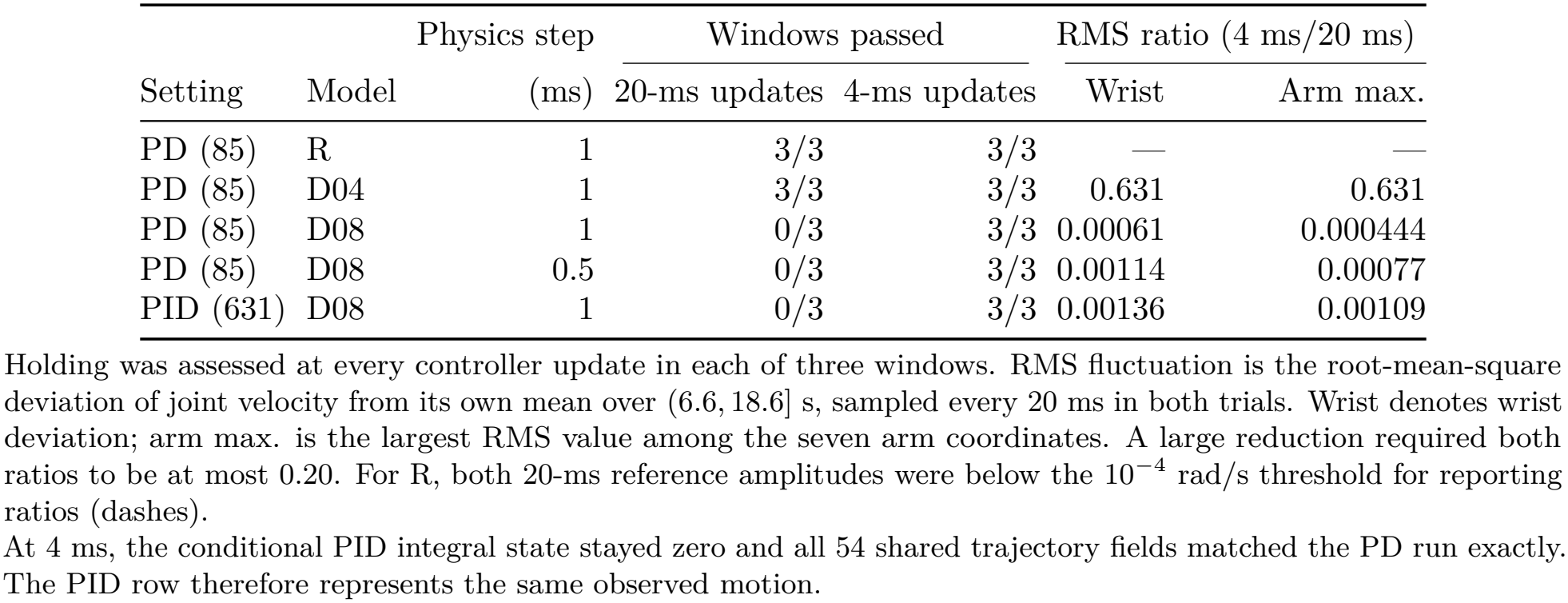
Post-hoc comparison of shortening the controller update period from 20 to 4 ms on holding and joint-speed fluctuations. Each row compares both update periods from the nominal start, with gains fixed. R is the native model; D04 and D08 include series elasticity and damping (Table 3).

**Table A2:** Can a map estimated from finite perturbations predict the full-state response? Each column represents one rigid-model controller setting. All four passed the 16 numerical and completeness checks, but none passed both amplitude comparisons or predicted the response to independent mixed-coordinate perturbations within the required bounds.

|  | Low PD | Middle reflex | High PID | Low reflex |
|---|---|---|---|---|
| Setting ID | 85 | 3749 | 4574 | 1870 |
| State dimension | 94 | 94 | 97 | 94 |
| Response map | *M*<sub>bound</sub> | *M*<sub>bound</sub> | *M*<sub>two</sub> | *M*<sub>two</sub> |
| *Agreement between maps at adjacent perturbation amplitudes* |  |  |  |  |
| 16 vs. 8 | pass | fail | fail | fail |
| 8 vs. 4 | fail | pass | fail | pass |
| *One-update prediction in independent mixed directions* |  |  |  |  |
| Eligible signed tests | 228 | 384 | 384 | 384 |
| Minimum relative error | 0.220 | 0.065 | 0.072 | 0.059 |
| Median relative error | 0.556 | 0.142 | 0.350 | 0.119 |
| Maximum relative error | 1.221 | 0.837 | 1.461 | 0.238 |
| *Spectral radius of the estimated map (descriptive)* |  |  |  |  |
| Minimum | 0.971546 | 0.994863 | 0.997028 | 0.996353 |
| Maximum | 0.971967 | 0.994875 | 0.998885 | 0.996410 |

Low, middle and high denote target postures. *M*<sub>two</sub> uses two-sided perturbations; *M*<sub>bound</sub> uses one-sided responses where activation bounds restrict the perturbation direction. The numbers 16, 8 and 4 multiply the reference perturbation widths. A pass required agreement within 5% in both matrix norms for the full state and both diagonal blocks; the outcomes were the same at solver levels L0–L3. Relative errors summarize the physical/controller block over eligible tests of the indicated map (Sect. A.2.7). The joint prediction criterion also tests the solver state. Spectral radii span solvers L0 and L3 at all three amplitudes. These finite-amplitude estimates did not establish local closed-loop stability.

**Table A3:** Directional stiffness of the fingertip at three equilibria in the rigid 34-muscle model. Muscle activation was fixed. Forces were applied in the world *xy* plane with vertical displacement free. Each row represents one nominal equilibrium.

|  |  | Stiffness (N/m) |  | Relative error (%) |  |
|---|---|---|---|---|---|
| Target | Controller (ID) | Minimum | Maximum | Method agreement | Held-out probes |
| Low | PD (85) | 2.751 | 59.690 | 0.000485 | 0.175 |
| Middle | Reflex (3749) | -19.573 | 39.925 | 0.000633 | 0.110 |
| High | PID (4574) | -6.248 | 6.735 | 0.001107 | 0.114 |

Minimum and maximum are the eigenvalues of the symmetric part of *K<sub>xy</sub>* across directions in the plane. Both are positive at the low target; opposite signs at the middle and high targets indicate an indefinite matrix. Method agreement compares the implicit-function estimate with the force-probe estimate. Held-out probes test displacement predictions in independent force directions. All three measurements passed the validation criteria, including recovery of known isotropic and anisotropic spring additions. The results describe static model mechanics at fixed activation.

**Table A4:**
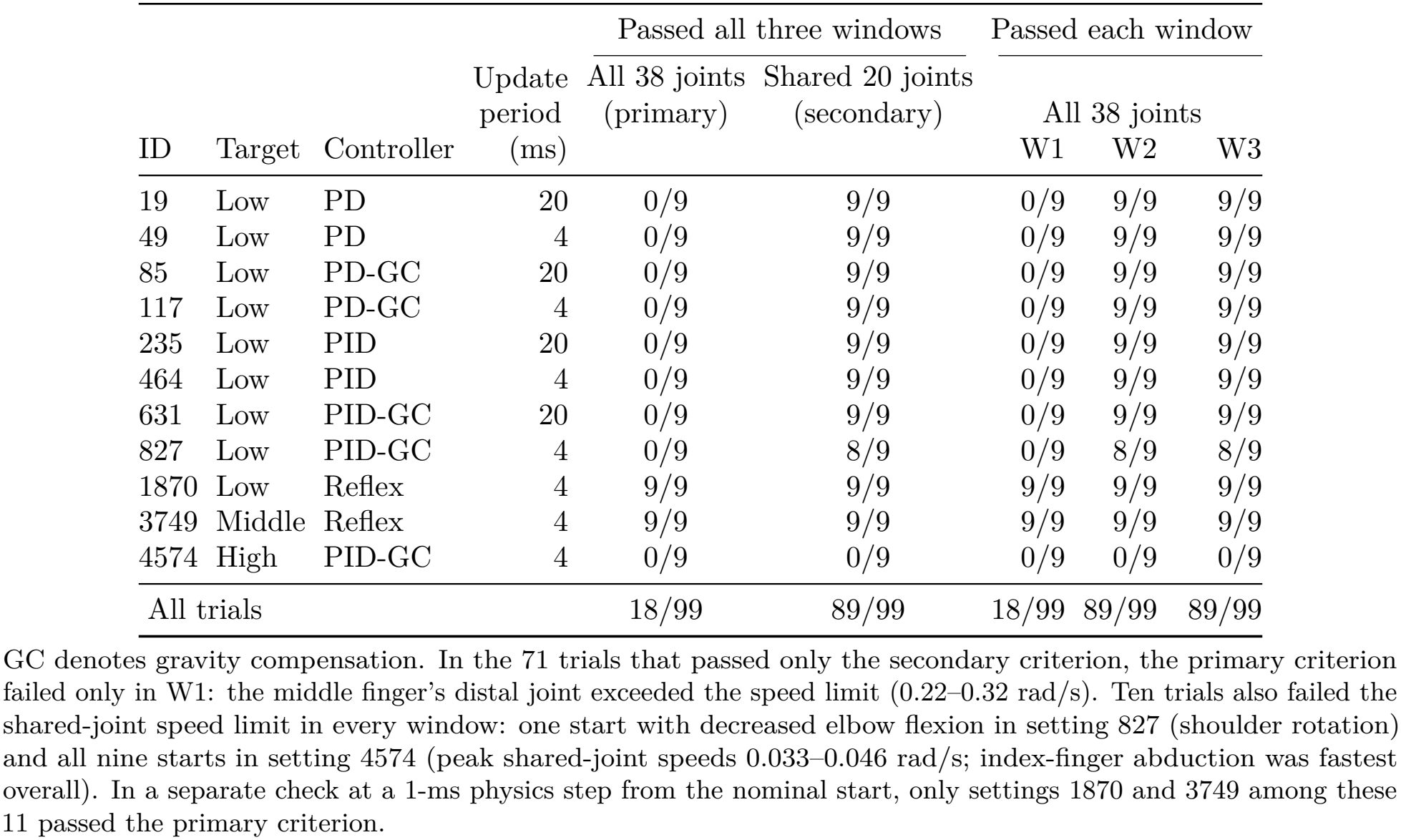
Holding after transfer of 11 settings from the 34-muscle model to the 63-muscle model without retuning. Each setting passed all three windows from all nine starts in the 34-muscle model. The table shows the same nine starts in the 63-muscle model (2-ms physics step). Both criteria require the same fingertip position and drift limits.

|  |  |  |  | Passed all three windows |  | Passed each window |  |  |
|---|---|---|---|---|---|---|---|---|
|  |  |  | Update period (ms) | All 38 joints (primary) | Shared 20 joints (secondary) | All 38 joints |  |  |
| ID | Target | Controller |  |  |  | W1 | W2 | W3 |
| 19 | Low | PD | 20 | 0/9 | 9/9 | 0/9 | 9/9 | 9/9 |
| 49 | Low | PD | 4 | 0/9 | 9/9 | 0/9 | 9/9 | 9/9 |
| 85 | Low | PD-GC | 20 | 0/9 | 9/9 | 0/9 | 9/9 | 9/9 |
| 117 | Low | PD-GC | 4 | 0/9 | 9/9 | 0/9 | 9/9 | 9/9 |
| 235 | Low | PID | 20 | 0/9 | 9/9 | 0/9 | 9/9 | 9/9 |
| 464 | Low | PID | 4 | 0/9 | 9/9 | 0/9 | 9/9 | 9/9 |
| 631 | Low | PID-GC | 20 | 0/9 | 9/9 | 0/9 | 9/9 | 9/9 |
| 827 | Low | PID-GC | 4 | 0/9 | 8/9 | 0/9 | 8/9 | 8/9 |
| 1870 | Low | Reflex | 4 | 9/9 | 9/9 | 9/9 | 9/9 | 9/9 |
| 3749 | Middle | Reflex | 4 | 9/9 | 9/9 | 9/9 | 9/9 | 9/9 |
| 4574 | High | PID-GC | 4 | 0/9 | 0/9 | 0/9 | 0/9 | 0/9 |
| All trials |  |  |  | 18/99 | 89/99 | 18/99 | 89/99 | 89/99 |

GC denotes gravity compensation. In the 71 trials that passed only the secondary criterion, the primary criterion failed only in W1: the middle finger's distal joint exceeded the speed limit (0.22–0.32 rad/s). Ten trials also failed the shared-joint speed limit in every window: one start with decreased elbow flexion in setting 827 (shoulder rotation) and all nine starts in setting 4574 (peak shared-joint speeds 0.033–0.046 rad/s; index-finger abduction was fastest overall). In a separate check at a 1-ms physics step from the nominal start, only settings 1870 and 3749 among these 11 passed the primary criterion.

